# Modeling the stepwise extension of recombination suppression on sex chromosomes and other supergenes through deleterious mutation sheltering

**DOI:** 10.1101/2021.05.17.444504

**Authors:** Paul Jay, Emilie Tezenas, Amandine Véber, Tatiana Giraud

## Abstract

Many organisms have sex chromosomes with large non-recombining regions that have expanded stepwise, generating “evolutionary strata” of differentiation. The reasons for this remain poorly understood, but the principal hypotheses proposed to date are based on antagonistic selection due to differences between sexes. However, it has proved difficult to obtain empirical evidence of a role for sexually antagonistic selection in extending recombination suppression, and antagonistic selection has been shown to be unlikely to account for the evolutionary strata observed on fungal mating-type chromosomes. There may, therefore, be other mechanisms involved in the extension of non-recombining regions. We show here, by mathematical modeling and stochastic simulation, that recombination suppression on sex chromosomes and around supergenes can expand in a stepwise manner under a wide range of parameter values simply because it shelters recessive deleterious mutations, which are ubiquitous in genomes. Permanently heterozygous alleles, such as the male-determining allele in XY systems, protect linked chromosomal inversions against the expression of their recessive mutation load, leading to the successive accumulation of inversions around these alleles without the need for antagonistic selection. Similar results were obtained with models assuming recombination-suppressing mechanisms other than chromosomal inversions, and for supergenes other than sex chromosomes, including those without XY-like asymmetry, such as fungal mating-type chromosomes. However, inversions capturing a permanently heterozygous allele were found to be less likely to spread when the mutation load was lower (e.g. under conditions of large effective population size, low mutation rates and high dominance coefficients). This may explain why sex chromosomes remain homomorphic in some organisms but are highly divergent in others. Here, we explicitly state and model a simple and testable hypothesis explaining the existence of stepwise extensions of recombination suppression on sex chromosomes, which can also be applied to mating-type chromosomes and supergenes in general.

## Introduction

Sex chromosomes, mating-type chromosomes and supergenes in general are widespread in nature and control striking polymorphisms, such as sexual dimorphism or color polymorphism, in many organisms, including humans (*1–5*). Supergenes display puzzling genomic architectures, defined by extensive regions lacking recombination, encompassing multiple genes (*3, 4*). Sex chromosomes, in particular, often appear to have evolved by stepwise extension of non-recombining regions, generating “evolutionary strata” of differentiation between haplotypes (*4, 6, 7*). It is generally thought that the reason for this gradual expansion of recombination suppression on sex chromosomes is that selection favors the linkage of sex-determining genes to sexually antagonistic loci [i.e. to alleles advantageous in one sex but disadvantageous in the other (*7–9*)]. However, theoretical concerns have been raised about this idea (*10*), and it has proved difficult to obtain empirical support for a role of antagonistic selection in the extension of recombination suppression (*6, 11, 12*). Furthermore, a gradual expansion of recombination suppression has been observed around many fungal mating-type loci (*5*) and at other supergenes, such as those controlling wing color in butterflies or social structure in ants (*13, 14*). Some type of antagonistic selection may exist between morphs determined by color or social supergenes, but antagonistic selection between fungal mating types is highly unlikely, given their life cycles and the results of genomic investigations (*15, 16*). For example, there has been a repeated stepwise extension of recombination suppression in anther-smut fungi despite the lack of a haploid phase consisting of cells of different mating types potentially expressing contrasted life history traits, leaving little room for antagonistic selection (*15, 16*). Alternative explanations have been put forward for the expansion of non-recombining regions on sex chromosomes (*11*), such as meiotic drive (*17*), genetic drift (*18*), deleterious-mutation sheltering (*19, 20*) and the neutral accumulation of sequence divergence (*21*), but the conditions in which such mechanisms could apply also appear to be restricted.

We develop here a general model for testing the idea that recombination suppression may extend stepwise around sex-determining or mating-type loci because it provides the fitness advantage of sheltering deleterious mutations segregating in nearby regions. Related hypotheses have been explored before, but in the restricted specific context of inbreeding (*19, 20*). We model here a more general hypothesis, based on the idea that genomes carry numerous deleterious recessive variants, as suggested by studies in a wide range of biological systems and by the pervasiveness of inbreeding depression in nature (*22–27*). We use mathematical modeling and stochastic simulations to test the hypothesis that permanently heterozygous alleles, such as male-determining alleles in XY systems, protect linked chromosomal inversions against the expression of their recessive mutation load, potentially leading to an accumulation of inversions around permanently heterozygous alleles, generating evolutionary strata.

The rationale behind this model is illustrated in Figure 1. Consider a diploid population carrying partially recessive, deleterious mutations. The combined effects of recombination, mutation, selection and drift result in individuals carrying different numbers of deleterious variants genome-wide and within particular genomic regions (Figure 1A). Chromosomal inversions may occur at any position and suppress recombination when heterozygous, thereby capturing a specific combination of deleterious variants (i.e. a haplotype). Inversions capturing fewer deleterious variants than the population average for the region concerned have a fitness advantage and should, therefore, increase in frequency. Such inversions are advantageous due to associative overdominance, i.e. the inversion itself is neutral but it captures a combination of alleles that is advantageous when heterozygous (*28, 29*). However, as the frequency of an inversion increases, homozygotes for this inversion become more common. Homozygotes are at a strong disadvantage due to the recessive deleterious variants carried by these inversions, and selection against homozygotes would therefore be expected to prevent such inversions from reaching high frequencies (Figure 1B).

**Figure 1.**
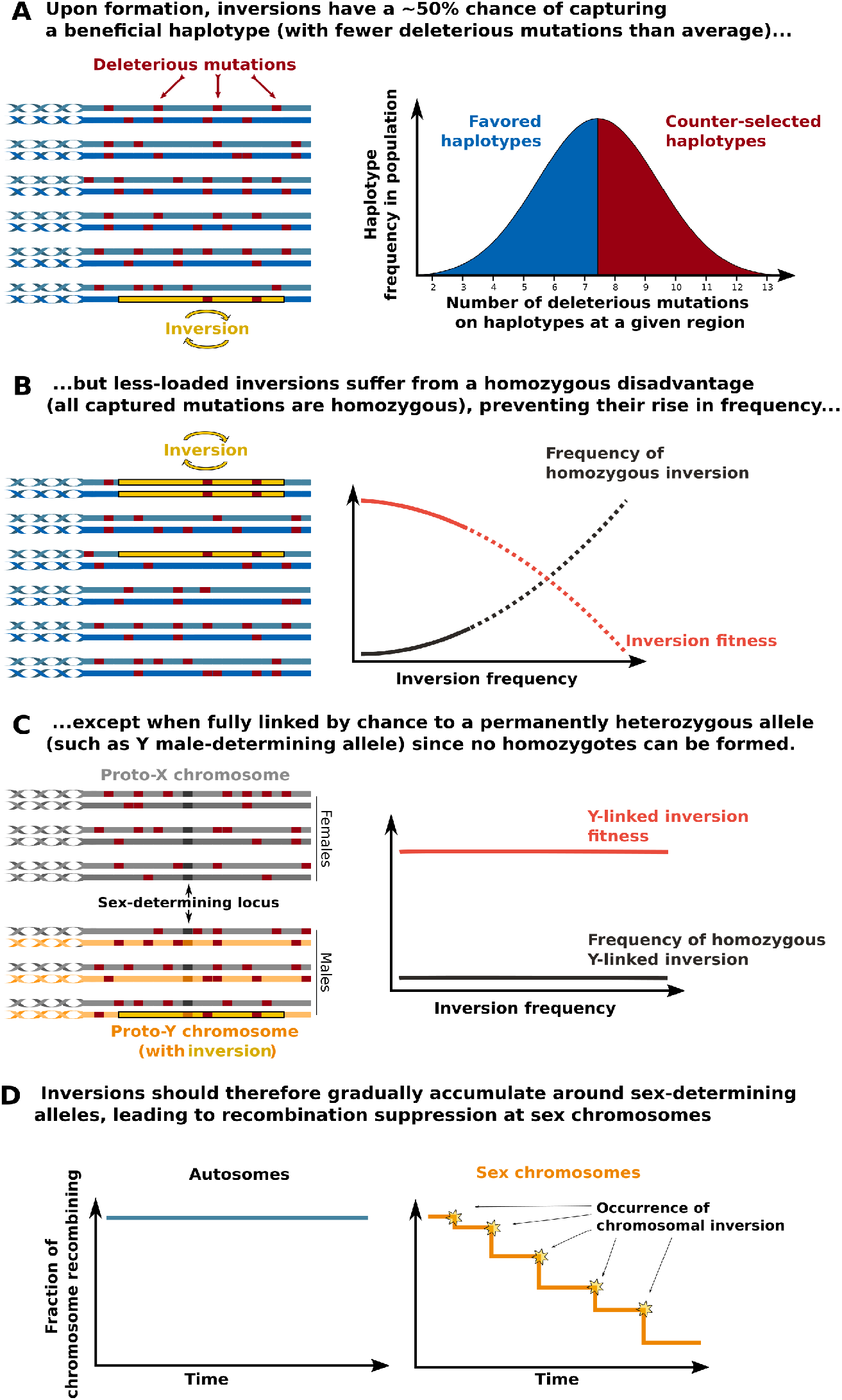
Schematic diagram of the model. **A**, Within any population and for any genomic region, diploid individuals (here represented by two homologous chromosomes) and haplotypes (i.e. combinations of mutations, here one chromosome) carry variable numbers of partially recessive, deleterious mutations. A substantial fraction (about 50%) of haplotypes have fewer deleterious mutations than the average and should be favored by selection. Chromosomal inversions therefore have a significant chance of capturing beneficial haplotypes. **B**, The increase in frequency of a beneficial inversion leads to an increase in the frequency of homozygotes for the inversion, which have a fitness disadvantage because they are homozygous for all the deleterious recessive mutations carried by the inversion. The fitness of the inversion therefore decreases with increasing inversion frequency, keeping the frequency of the inversion low. **C**, Permanent heterozygosity at a Y-chromosome sex-determining allele protects linked inversions from this disadvantage of homozygosity. Hence, beneficial inversions (carrying fewer deleterious mutations than average) should spread and become fixed in the population of Y chromosomes.**D**, Unlike those on autosomes, inversions capturing the Y sex-determining allele never suffer from homozygous disadvantage and should therefore accumulate on proto-Y chromosomes, leading to a suppression of recombination between the X and Y chromosomes. A similar mechanism should operate for any recombination suppressor acting in *cis* (not just chromosomal inversions) and any locus with permanently heterozygous alleles.

Now, consider an inversion that, by chance, captures a permanently heterozygous allele, such as the male-determining allele in an XY system. If this Y-linked inversion captures fewer deleterious variants than the population average, it should increase in frequency without ever suffering the deleterious consequences of having its load expressed. The recessive deleterious mutations captured by the sex-linked inversion are, indeed, fully associated with the permanently heterozygous, maledetermining allele, and will, therefore, never occur as homozygotes. Unlike autosomal inversions, Y-linked inversions retain their fitness advantage with increasing frequency (Figure 1C). Hence, less heavily loaded Y-linked inversions would be expected to spread, becoming fixed in the population of Y chromosomes, resulting in a suppression of recombination between the X and Y chromosomes in the region covered by the inversion.

The successive fixation of additional inversions linked to this Y-fixed inversion should cause the non-recombining region to expand further, by the same process, thereby leading to the formation of a chromosome with a large non-recombining region around the sex-determining locus —i.e. a sex chromosome with evolutionary strata of differentiation (Figure 1D). We use the term “chromosomal inversions” here for simplicity, but the proposed mechanism would hold for any mechanism suppressing recombination, and we model several types of recombination suppressor. The proposed mechanism would be expected to apply to any genomic region with a permanently or near-permanently heterozygous locus, such as mating-type loci in fungi or supergenes in many organisms. The accumulation of deleterious mutations following recombination suppression has been extensively studied (*30, 31, 31–33*), but we investigate here its converse: that deleterious mutations could be a cause, and not only a consequence, of recombination suppression.

## Results

We explored this general hypothesis of stepwise recombination suppression around permanently heterozygous alleles with infinite population deterministic models and individual-based simulations. Very little is known about the dynamics of deleterious mutations in genomic regions under particular recombination regimes, such as those generated by polymorphic inversions. We therefore had to use simulations to explore realistic scenarios involving various levels of drift. In both the infinite population model and individual-based simulations, we modeled diploid populations with only partially recessive deleterious mutations, occurring at a rate *u*, with heterozygous and homozygous individuals suffering from *1-hs* and *1-s* reductions in fitness, respectively, and with multiplicative effects of mutations on fitness. We first considered that all mutations occurring in genomes had the same dominance (*h*) and selection (*s*) coefficients, and then relaxed this hypothesis. Individuals were considered to have two pairs of chromosomes, one of which harbored a locus with at least one allele permanently or almost permanently heterozygous (see Methods for details). Several situations were considered, mimicking those encountered in XY sex-determination systems, in fungal mating-type systems and in overdominant supergenes. We analyzed the evolution of recombination modifiers suppressing recombination across the fragment in which they reside (i.e. *cis*-modifiers), either exclusively in heterozygotes (mimicking for example chromosomal inversions), or in both heterozygotes and homozygotes (e.g. histone modifications). Each of these recombination modifiers, which was assumed to be neutral in itself, appeared in a single haplotype, and was, thus, in linkage disequilibrium with a specific set of mutations, such that its fitness was exclusively dependent on the number of deleterious alleles within the segment captured. We first compared the dynamics of inversion-mimicking mutations in an autosome with those capturing a male-determining allele in an XY system (males are XY and females are XX, the male-determining allele being permanently heterozygous), and we then considered other types of recombination modifiers and heterozygosity rules.

As noted by several authors (*28, 34, 35*), inversions capturing fewer deleterious mutations than the population average should increase in frequency. In infinite populations, the number of mutations harbored by individuals within a genomic segment of size *n* follows a binomial distribution of parameters *n* and *q*, with *q* the mean frequency of mutations at mutation-selection equilibrium (Figure 2a and Figure S1). On average, individuals harbor *nq* mutations on each of their chromosomal segments of size *n*. Under realistic parameter values, the vast majority of large chromosomal regions therefore carry several deleterious mutations. For instance, considering *s*=0.001, *h*=0.1, *u*=1e-09 and *n*=2Mb, more than 99.999% of chromosomal fragments carry a least one mutation, the mean number of mutations being *nq* = 20 (Figure S1). If *m* is the number of recessive deleterious mutations captured by a given inversion, the mean fitness of individuals homozygous for the inversion (W_II_), heterozygous for the inversion (W_NI_) or lacking the inversion (W_NN_) in an infinite population can be easily expressed as a function of *n, q, h, s*, and *m* [see Methods; (*28, 34*)]. Once formed, inversions should increase in frequency if heterozygotes for the inversion are fitter than homozygotes without the inversion (W_NI_>W_NN_), which is the case if the inversion carries fewer mutations than the population average [*m* < *nq;* (*28*)].

**Figure 2.**
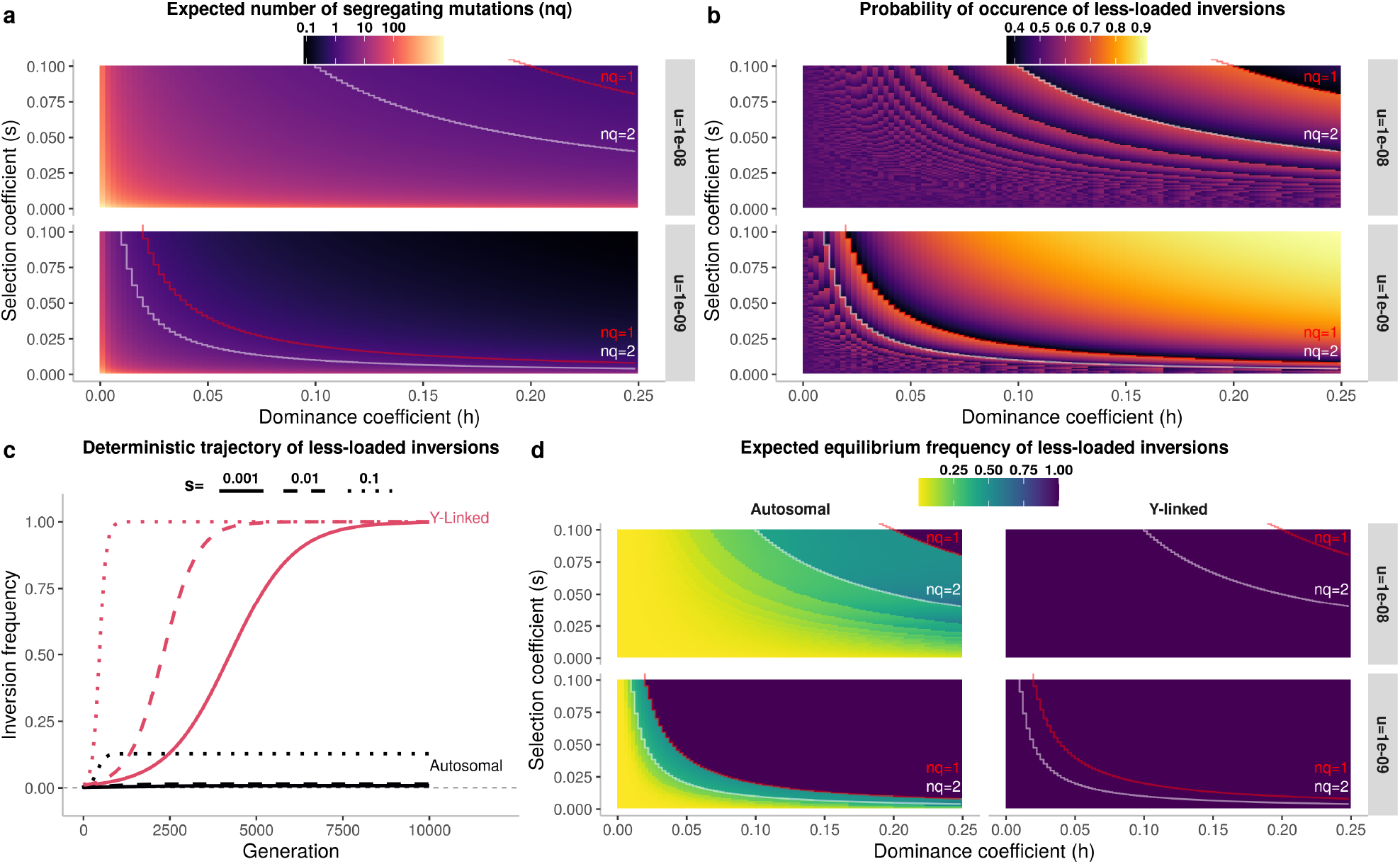
In infinite populations, all less-loaded chromosomal inversions become fixed on the Y sex chromosome, whereas most remain at a low frequency on the autosome. **a**, Expected number of segregating recessive deleterious mutations within a segment of *n*=2 Mb as a function of the dominance coefficient (*h*), selection coefficient (*s*) and mutation rate (*u*). Mutations were considered to be at mutation-selection equilibrium frequency (see methods). Parameters resulting in one and two expected mutations are highlighted by red and white lines, respectively, and are also shown on panels b and d. **b**, Probability of occurrence of a less-loaded inversion covering a segment of *n*=2 Mb as a function of the dominance coefficient (*h*), selection coefficient (*s*) and mutation rate (*u*). Less-loaded inversions are defined as those for which *m*<*nq*, where *m* is the number of mutations in the inversion (i.e., fewer than the average for the population). The number of mutations in inversions can take only integer values (*m*=0, 1, 2, etc.), and the transition between these values affects the probability of occurrence of less-loaded inversions, yielding the non-continuous graphs in panels b and d. **c**, Deterministic change in inversion frequency, for the case of 2 Mb inversions capturing the male-determining allele on the Y-chromosome or inversions on an autosome. For clarity, the frequency displayed for Y-linked inversions is the frequency of inversions in the population of Y chromosomes. The figure illustrates the case of inversions carrying a number of mutations 5% lower than the population average (*m*=[0.95\**nq*]), mutations being at their mutation-selection equilibrium frequency with *u*=10^-8^ and *h*=0.1. Figures S4-8 illustrate the fate of inversions with intermediate degrees of linkage to the male-determining allele, with various numbers of mutations relative to the population average, or inversions linked to a locus with variable heterozygosity rules or in the X chromosome population. **d**, Expected equilibrium frequency of less-loaded 2 Mb inversions (i.e. with *m*=0,1,2, …, nq) capturing the male-determining allele, or on an autosome, weighted by their probability of occurrence (Methods, equation 5, 6, and 7). This panel extends the result from panel c for a range of dominance coefficients (*h*), selection coefficients (*s*) and mutation rates (*u*). As above, the frequency displayed for Y-linked inversions is the frequency of inversions on the population of Y chromosomes. Above the line *nq*=1, the mean number of segregating mutations in a given inversion (of length *n*) is less than 1, indicating that all less-loaded inversions are mutation-free. Such inversions can also become fixed on autosomes, leading to an expected equilibrium frequency of 1 for both autosomes and the Y chromosome. Figure S9 shows the overall frequencies of all inversions, whether less or more loaded than average, and over a larger range of parameters.

Repeated sampling from binomial distributions with a wide range of parameters showed that more than half the inversions occurring in genomes captured fewer deleterious mutations than the population average (figure 2b; hereafter referred to as “less-loaded inversions”). Indeed, the distribution of mutation number across individuals is almost symmetric when *n*q* is high (as the binomial distribution converges to a normal distribution; e.g. Figure S1), but, when *n*q* is low, the distribution is zero-inflated, increasing the probability of inversions being less-loaded. Therefore, a substantial fraction of inversions occurring in genomes are beneficial when they form (i.e. when rare enough not to occur as homozygotes). For instance, with *h* values ranging from 0 to 0.5 and *s* values ranging from 0.001 to 0.25, between 36% and 98% (mean=70%) of the 2 Mb inversions occurring in the genome are beneficial, carrying fewer recessive deleterious mutations than average. Simulations in finite populations of different sizes confirmed that most inversions (mean=66% in the range of parameter values studied) had a fitness advantage upon formation. The simulations also showed that inversions could be favored if they captured mutations that were rarer than average (Figures S2-3).

The deterministic increase in frequency of an inversion (I) on an autosome or capturing an XY-like sex-determining locus can easily be determined with a two-locus two-allele model. For instance, the change in frequency of an inversion on the proto-Y chromosome can be expressed as:

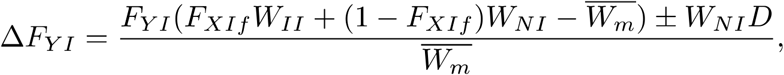

where *F_XIf_* is the frequency of the inversion on the proto-X chromosome in females, *r* is the rate of recombination between the inversion and the sex-determining locus, *D* is their linkage disequilibrium and 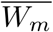 is mean male fitness (Figure 2c; see Methods and Appendix for details). Based on this model, and initially assuming that inverted and non-inverted segments no longer accumulate deleterious mutations after their formation (i.e. W_II_, W_NI_ and W_*NN*_ are fixed parameters), we simulated the evolutionary trajectory of inversions on autosomes and of inversions capturing the maledetermining allele on the Y chromosome under a wide range of parameter values. We found that the frequency of less-loaded inversions tended to remain low in autosomes, whereas these inversions became fixed in the population of Y chromosomes (Figure 2c), as expected according to our hypothesis.

We therefore expressed the equilibrium frequency of inversions (*F_equ,m_*, Figure 2d) as a function of *m*, the number of deleterious mutations carried by the inversion, to calculate the threshold values at which autosomal and Y-linked inversions could be maintained or fixed (see appendix). Assuming that *q*<<1 and *s*<<1, we found that inversions on autosomes become fixed when *m*<*qhn*/(1-*h*), whereas they should stabilize at a frequency of 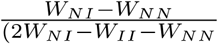, the equilibrium frequency of an overdominant locus, when *qhn*/(1-*h*)<*m*<*nq* (see Methods). Inversions on autosomes capturing more than *qhn*/(1-*h*) recessive deleterious mutations do, indeed, suffer from a homozygote disadvantage (WNI > WII, figure S1), preventing them from reaching high frequencies (Figure 2c). This is the case for most autosomal inversions under realistic parameter values (e.g. *m*>*qhn*/(1-*h*) for more than 99.99 % of inversions when *n*=2 Mb, *u*=1e-09, *h*=0.1 and *s*=0.001, Figure S1). By contrast, permanently heterozygous alleles protect inversions from this homozygote disadvantage, allowing these inversions to become fixed. All inversions capturing the male-determining allele become, therefore, fixed if they carry fewer than average mutations (i.e. *m*<*nq*). Thus, contrary to the argument proposed in a previous study that only mutation-free inversions can become fixed (*35*), we found that inversions can carry deleterious mutations and nevertheless become fixed in the population, provided that they carry fewer than *qhn*/(1-*h*) mutations if they are located on autosomes and fewer than *nq* mutations if they capture a permanently heterozygous allele (on the Y chromosome, for example).

The expected equilibrium frequency of all inversions occurring in a given genomic region can be expressed as :

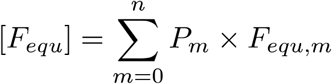

with *P_m_* being the probability of occurrence of inversions with *m* mutations and *F_equ,m_* the equilibrium frequency of inversions with *m* mutations (Figure 2d; see Methods). For permanently heterozygous inversions, we, thus, have:

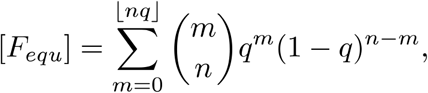

and for autosomal inversions:,

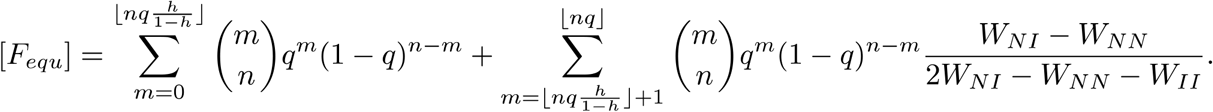

With a realistic range of parameters, inversions are much more likely to become fixed if they capture the male-determining allele on the Y chromosome than if they are unlinked to this allele (e.g. on an autosome; Figure 2c-d and Figures S4-9). For instance, with *u*=1e-09, *h*=0.1, *s*=0.001, and *n*=2 Mb, 47% of inversions occurring on the Y chromosome would be expected to become fixed, versus only 0.000045% of inversions on autosomes.

If deleterious mutations continue to arise after the formation of the inversion, the dynamics of the inversion become harder to predict deterministically. In infinite populations, the accumulation of mutations after the formation of the inversion can be approximated by 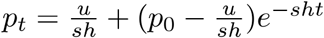 where *P_t_* is the number of mutations in the inversion at time *t* (*28, 34*). This assumes that inverted segments evolve under mutation-selection dynamics with recombination. However, such assumptions do not hold in many situations. Indeed, low-frequency inversions in finite populations should almost never occur as homozygotes and should therefore evolve with almost no recombination. This is also true for inversions capturing a permanently heterozygous allele, which never undergoes recombination. We confirmed, by simulation, that the above approximations for an infinite population depart strongly from the situation observed in finite populations (e.g. Figure S13). In finite populations, low-frequency or permanently heterozygous inversions tend to evolve under a Muller’s ratchet-like dynamic, with the mean fitness of inversions decreasing in a stepwise manner due to the sequential loss, by drift, of the inverted haplotypes with the lowest mutational load. Following their formation, autosomal and Y-linked inversions tend to accumulate more mutations than the average for the population, contrasting with predictions for infinite populations (Figure S13). Only inverted segments reaching relatively high frequencies in autosomes eventually recombine when homozygous, and their dynamics of mutation accumulation therefore involve a mixture of a Muller’s ratchet-like regime (when rare, at the start of their spread) and a mutation-selection-drift regime with recombination (when they reach intermediate frequencies). Little is known about the transition between these regimes (*36, 37*). We therefore used individual-based simulations to study the fate of inversions accumulating deleterious mutations. This also made it possible to relax the previous assumption (*35*) that the time to inversion fixation is much shorter than the time taken for the inversions to accumulate deleterious mutations. Individual-based simulations also made it possible to relax the assumption that all genomic positions are independent.

Simulations of the spread of inversions in finite populations with recurrent mutation confirmed the tendency identified in the deterministic model without mutation accumulation: over most of the parameter space, inversions are much more likely to spread if they capture the sex-determining allele on the Y chromosome than if they are located on autosomes (Figure 3 and Figures S10-16). Many autosomal inversions carrying a mutation load segregated for hundreds of generations. For example, with *N*=1000, *s*=0.01, *h*=0.1 and *u*=1e^-08^, 73 of 10,000 inversions of 2 Mb in length continued to segregate after 500 generations. However, all these autosomal inversions were lost at the end of the simulations (i.e. after 10,000 generations, Figure 3a). These inversions initially spread because, by chance, they had a lower-than-average mutation load, but their homozygote disadvantage prevented them from reaching fixation. They were eventually lost because they accumulated further mutations relative to non-inverted segments (Figure 3b and S13).

**Figure 3.**
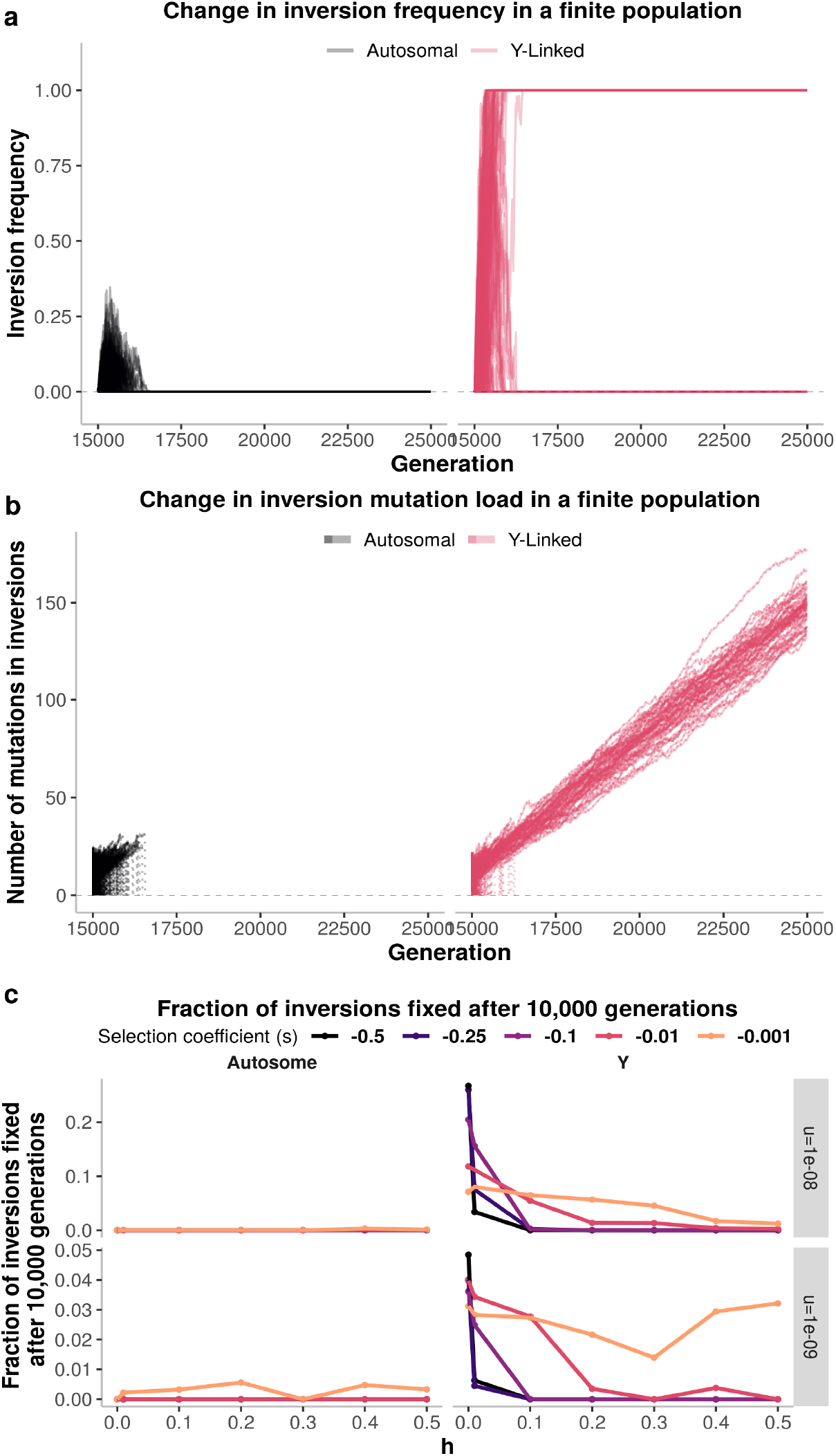
In finite populations, a substantial fraction of less-loaded chromosomal inversions on the Y sex chromosome become fixed, whereas all less-loaded inversion on autosomes are lost. **a**, Change in inversion frequency in stochastic simulations of 1000 individuals experiencing recessive deleterious mutations at a rate u=1e-08, all mutations having the same dominance and selection coefficients (here, *h*=0.1 and *s*=0.01). The figure displays the frequency of 10,000 independent inversions on each of an autosome and a proto-Y chromosome, with each line representing a specific inversion (i.e. a simulation). The evolutionary trajectories of inversions of different sizes, in larger populations, on the X-chromosome or with other parameter combinations are displayed in figures S10-14. **b**, Change in the mean number of mutations carried by the inverted segments in each of the 10,000 simulations. Each line represents one simulation. A zoom-view of the autosomal inversion dynamics is shown in Figure S13. **c**, Proportions, in stochastic simulations, of 2 Mb inversions that were fixed after 10,000 generations, for different parameter value combinations. This extends the result from panel **a** to other values of dominance coefficient (*h*), selection coefficient (*s*) and mutation rate (*u*). For each parameter combination, we simulated 10,000 inversions of 2 Mb capturing a random genomic fragment. Only inversions not lost after 20 generations are considered here. Results for inversions of different sizes are shown in Figure S15 and results for different population sizes are shown in Figure S16.

By contrast, substantial fractions of less-loaded inversions capturing the permanently heterozygous sex-determining allele on the Y chromosome spread until they became fixed in the Y chromosome population; this was the case even for inversions that were not mutation-free (Figure 3 and Figure S10-16). For example, for *s*=0.01, *h*=0.1, *u*=1e^-08^ and *N*=1000, 49 of the 10,000 Y-linked inversions (2 Mb) became fixed in the Y chromosome population, whereas all inversions on the autosome were lost. New mutations occurred on Y-linked inversions, but they did not accumulate rapidly enough to prevent these 49 inversions from spreading and reaching fixation (Figure 3a-b and Figure S13). Permanent heterozygosity effectively results in directional selection for the less-loaded inversions, and not only for those free from mutations (Figure 3), leading to rapid fixation before the accumulation of too many new deleterious mutations.

Similar results were obtained when: i) two or more permanently heterozygous alleles segregated at a mating incompatibility locus, modeling plant self-incompatibility or fungal mating-type systems [Figure S5 and S14; (*5, 38*)]; ii) the alleles were not permanently heterozygous, but were strongly overdominant, thus occurring mostly in the heterozygous state, as for several supergenes controlling color polymorphism [Figure S8; (*13, 39, 40*)]; and iii) the inversion was in strong but incomplete linkage with the permanently heterozygous allele (e.g. 0.1 cM away from the allele, Figure S4-8). Our model can thus account for the existence of inversions very close, but not fully linked to a mating-type locus, as reported in the chestnut blast fungus (*41, 42*). Similar results were also obtained when we considered recombination modifiers suppressing recombination even when homozygous, as for histone modifications or methylation (*43*), rather than solely when heterozygous, as for inversions (Figure S14). Similar results were also observed if we assumed that the fitness effects of mutations occurring across the genome were drawn from a gamma distribution (i.e. the mutations segregating in populations had different fitness effects; Figure S17).

We, thus, found that neutral recombination modifiers spread in a very large range of conditions, if they captured permanently heterozygous alleles (Figure 3C and Figure S15-16). However, as expected, for the inversions to benefit from a heterozygote advantage by associative overdominance, deleterious mutations segregating in the genome had to be partially or completely recessive (h<0.5; Figure 3c and Figures S15-16). Furthermore, as for any variant, even beneficial inversions benefiting from a selective advantage could be lost by drift, the probability of such loss depending on population size and the relative fitness of the inversion, in terms of the number and type of mutations initially captured relative to the average for the population (Figures 3 and S15-16). Inversions occurring in regions in which large numbers of mutations segregate are more likely than those in mutation-poor regions to capture many fewer mutations than average, and therefore to have a stronger relative advantage. In other words, inversions in mutation-dense regions have a wider fitness distribution. The probability of Y-linked inversion fixation, thus, increases with increasing inversion size, mutation recessiveness and mutation rate (Figure 3c and Figures S15-16). As expected (*44*), we found that, with increasing population size, beneficial inversions took more generations to spread to fixation (Figure S11-12). This longer time favored the further accumulation of deleterious mutations in inversions, lowering their fitness and, in some cases, preventing their fixation. The probability of inversion fixation in the Y chromosome population was, therefore, inversely correlated with population size, but was always much higher than that of fixation on autosomes (Figure S15-16).

Inversions would, therefore, be expected to accumulate successively around permanently heterozygous alleles across evolutionary time, under a large range of conditions, leading, for example, to the formation of non-recombining sex chromosomes with a typical pattern of evolutionary strata. We studied the formation of such strata by the sequential occurrence of multiple inversions, by simulating the evolution of large chromosomes experiencing the occurrence of multiple chromosomal inversions that can overlap with each other, under parameter values typical of those observed in mammals (Figures 4, 5 and S19). We simulated, over 100,000 generations, populations of N=10,000 or N=1,000 individuals, carrying two 100 Mb chromosomes, one of which harbored a mammalian-type sex-determining locus (XY males and XX females). Individuals experienced only deleterious or nearly-neutral mutations, with mutation rates and fitness effects similar to those observed in humans (fitness effect being drawn from a gamma distribution). The dominance coefficient of each mutation was chosen uniformly at random from a wide set of values (see Methods for details). At the start of each simulation, we randomly sampled *k* genomic positions that could be used as inversion breakpoints, with *k* being 10, 100, 1000 or 10,000. These genomic positions represent inversion hotspots, such as those that can be generated by repeats in genomes (*45*). In each generation, we introduced *j* inversions, *j* being sampled from a Poisson distribution. Simulation times were limited by using inversion rates such that one inversion, on average, occurred in each generation, throughout the whole population. The two breakpoints of each inversion were chosen, at random, from the k positions. It was, therefore, possible for two independent inversions to appear at the same position, allowing, in particular, the reversion of an inversion to its ancestral orientation, thereby restoring recombination. Reversion may be favored, for example, by an inversion accumulating enough deleterious mutations after its fixation on the Y chromosome to carry a larger number of mutations than the population average (*46*). We assumed that inversions partially overlapped by another inversion or that captured a smaller inversion could not be reversed, i.e., recombination could not be restored in such situations even if subsequent inversions reused the same breakpoints. Indeed, reversions of partially overlapped inversions do not restore ancestral arrangements but instead result in complex reshufflings of gene order and orientation and are therefore unlikely to restore recombination. We ran 10 simulations for each set of parameters.

**Figure 4.**
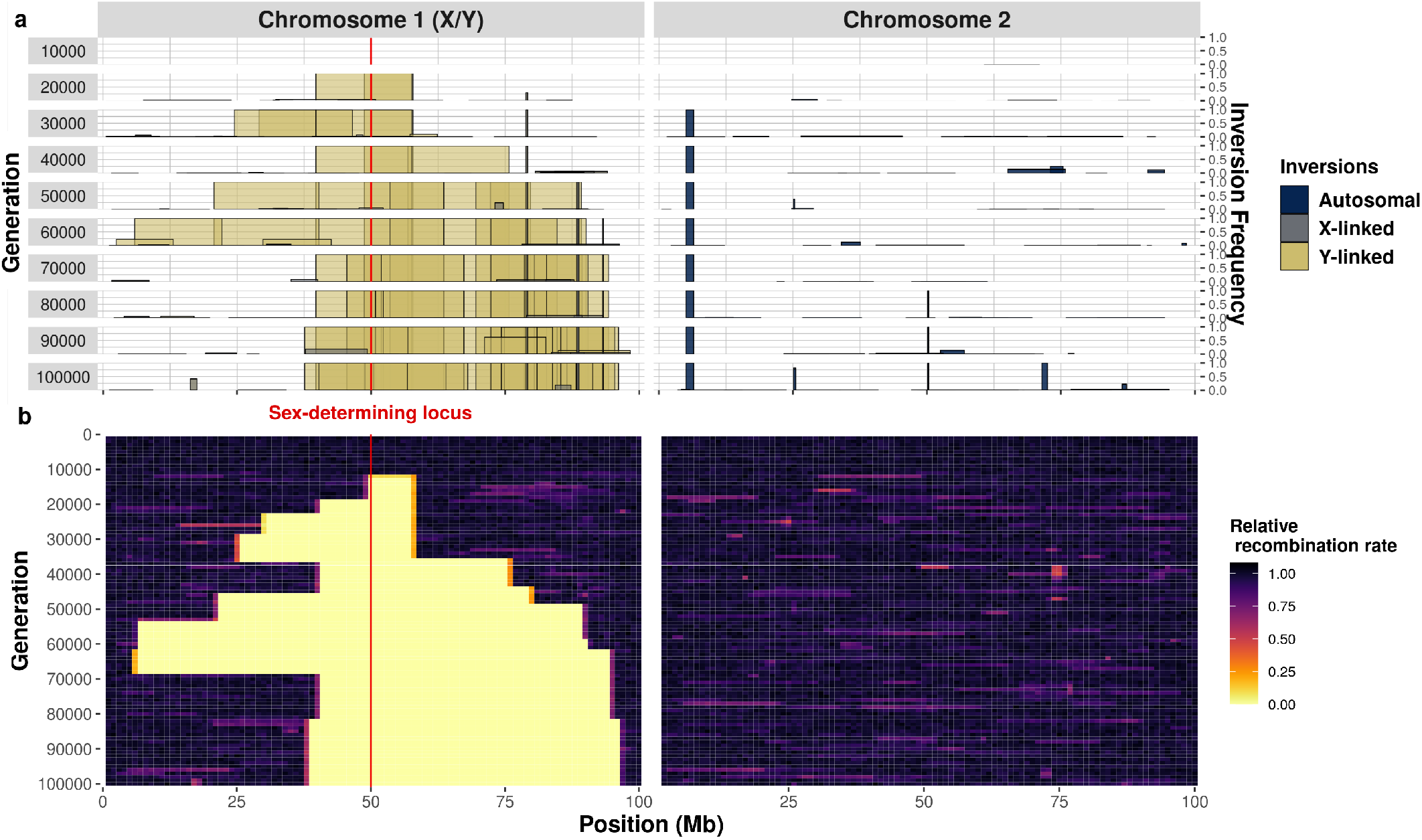
Successive accumulation of inversions around a male-determining allele in an XY system, leading to the formation of non-recombining sex chromosomes. A simulation of *N*=1000 individuals, each with two pairs of 100 Mb chromosomes, over 100,000 generations. Chromosome 1 harbors an X/Y sex-determining locus at 50 Mb (individuals are XX or XY). In each generation, one inversion appears, on average, in the whole population, in an individual sampled uniformly at random, with the two recombination breakpoints sampled uniformly at random from *k*=100 potential breakpoints. **a,** Overview of chromosomal inversion frequency and position for 10 different generations. The width of the box represents the position of the inversion and the height of the box indicates inversion frequency. Inversions appearing on the Y chromosome are depicted in yellow, those appearing on the X chromosomes are depicted in gray. The colors are not entirely opaque, so that regions with overlapping inversions appear darker. Previously fixed inversions may be lost due to the occurrence of beneficial reversions and selection. **b,** Changes in the relative rate of recombination over the entire course of the simulation. The numbers of recombination events occurring at each position (binned in 1 Mb windows) are recorded at the formation of each offspring, across all homologous chromosomes in the population. Only recombination events between the X and the Y chromosomes are shown for chromosome 1 (i.e., recombination events between the two X chromosomes in females are not shown). Unlike chromosome 1, chromosome 2 harbors no permanently heterozygous alleles. All inversions on this chromosome suffer from homozygote disadvantage and very few inversions therefore become fixed on chromosome 2. See Figure S19 for a simulation with *N*=10,000 individuals.

**Figure 5.**
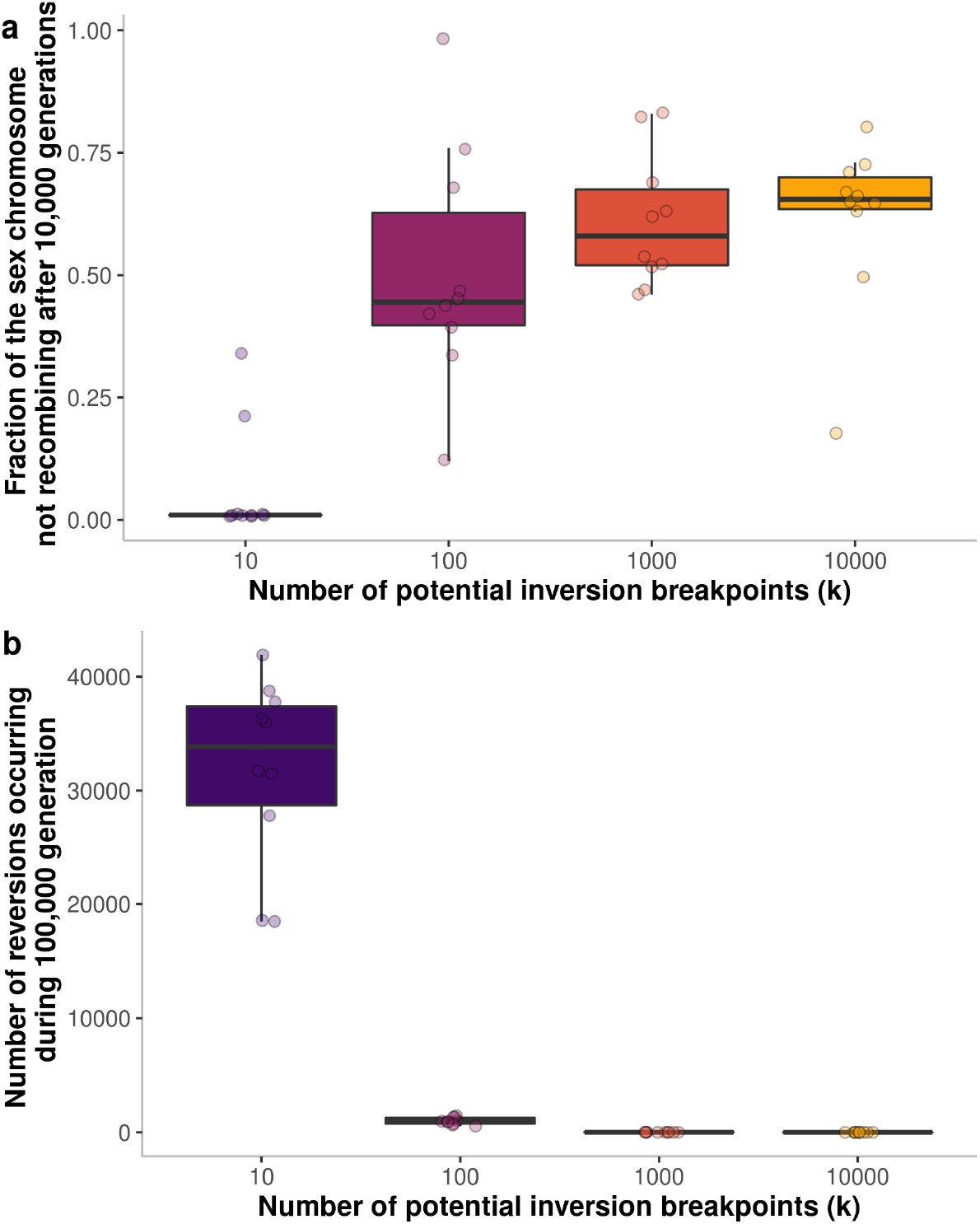
Effect of the number of potential inversion breakpoints on the evolution of recombination suppression. For each number of breakpoints, 10 simulations were conducted. Each dot represents the result of a simulation with *N*=1000. **a,** Fraction of the length of the Y sex chromosome not recombining after 100,000 generations. **b,** Number of reversions occurring over the course of the 100,000 generations. Boxplot elements: central line: median, box limits: 25th and 75th percentiles, whiskers: 1.5x interquartile range.

In all simulations assuming relatively large numbers of potential inversion breakpoints (k=100, 1000, 10,000), the Y chromosome progressively stopped recombining with the X chromosome as it accumulated inversions fully linked to the male sex-determining allele (Figures 4, 5 and S19). After the occurrence of an initial inversion capturing the male sex-determining allele, multiple inversions partially overlapping this inversion or other Y-fixed inversions were selected, thereby generating a growing chaos of overlapping chromosomal rearrangements. The non-recombining region, thus, extended around the sex-determining locus in a stepwise manner, perfectly reflecting the evolution of sex chromosomes and other supergenes with evolutionary strata (Figure 4). Some events in the gradual extension of recombination suppression were reversed, due to the occasional occurrence of beneficial reversions (Figures 4 and 5). The accumulation of overlapping inversions was, however, more rapid than the occurrence of beneficial reversions, leading to a progressive extension of the non-recombining region (Figure 4 and 5). By contrast, when we assumed a smaller number of potential breakpoints (*k* = 10), recombination suppression between sex chromosomes evolved in only two of the 10 simulations, and across only a small genomic region (Figure 5), owing to the more frequent occurrence of beneficial reversions.

## Discussion

Our results show that recombination suppression on sex chromosomes and other supergenes can evolve simply because genomes harbor many partially recessive, deleterious variants. Our model for the evolution of sex chromosomes, and supergenes in general, is based on simple and widespread phenomena: i) inversions (or any recombination suppressor) can be favored solely because they contain fewer deleterious mutations than the population average, a situation applying to a substantial fraction of the inversions formed; ii) such inversions tend to display overdominance: they are beneficial in the heterozygous state but suffer from a homozygote disadvantage, which prevents them from reaching high frequencies and becoming fixed on autosomes; iii) when, by chance, inversions capture a permanently heterozygous allele, they do not suffer from this homozygote disadvantage and are therefore able to spread until they are fully associated with the permanently heterozygous allele (e.g. they become fixed in the Y chromosome population). These three phenomena have been reported independently in several studies, but, to our knowledge, never in interaction [see references (*28, 35*) for i, (*29, 47*) for ii, and (*5, 19*) for iii]. The combined influence of the mechanisms related to ii and iii has been shown to promote sex chromosome-autosome fusion in highly inbred populations (*20*). We show here that such mechanisms can readily lead to the stepwise extension of the non-recombining region on sex chromosomes themselves, without the need for inbreeding. Moreover, unlike previous studies [e.g. (*18, 35, 48*)], we show that the higher probability of inversion fixation on Y chromosomes is not restricted to mutation-free inversions, but applies to any inversion capturing fewer deleterious mutations than the average.

We did not include possible decreases in fertility due to segregation issues arising during meiosis as a result of inversions in our model (*49*); such issues would, clearly, decrease the probability of inversion fixation as a function of the fitness cost, as for any other model explaining the evolution of the stepwise extension of recombination suppression involving inversions. However, our model holds for any mechanism of recombination suppression (it is not limited to inversions) and the other possible mechanisms [such as methylation (*43*)] would not be expected to decrease fertility. Given the lack of knowledge about inversion rates in natural conditions and the computation challenge represented by the simulation of complex patterns of recombination, we used reasonably high inversion rates in our sexchromosome evolution simulations, making it possible to observe the stepwise extension of a nonrecombining region within 100,000 generations. A recent study suggested that inversions may not be too rare and that inversion breakpoints may be widely distributed throughout the genome (*45*). The use of different inversion rates might result in much shorter or much longer times for the stepwise extension of the non-recombining region, but should not change the final outcome. In nature, the stepwise extension of the non-recombining region between sex or mating-type chromosomes often occurs over time scales of the order of tens of millions of generations [e.g. about 250 M years in human (*50*)], suggesting that the fixation of inversions may be a relatively rare event.

On autosomes, inversions maintained at low frequencies because of their homozygote disadvantage tend to be lost rapidly because they accumulate further deleterious mutations. On the Y chromosome, contrary to previous suggestions (*28*), we show that the accumulation of further deleterious mutations following the formation of an inversion is generally too slow to prevent the fixation of less-loaded inversions. Deleterious mutations nevertheless accumulate following fixation of the inversion, leading to Y-linked inversion degeneration, as observed on many non-recombining sex chromosomes (*30*). This may lead to selection for reversion of the inversion, thereby restoring recombination (*46*). The question as to whether this can actually occur is common to all mechanisms explaining evolutionary strata, including sexual antagonism.

We found inversion reversion to be rare, unless there were no more than 10 inversion breakpoints over the two 100 Mb chromosomes. Indeed, for reversions to invade the population and restore recombination, the following conditions must be met: i) sufficient deleterious mutations must accumulate in the initial inversion to render reversion beneficial and ii) a partially overlapping inversion should not have invaded the population before the occurrence of a beneficial reversion. Indeed, partially overlapping inversions result in a complex reshuffling of gene order and orientation, preventing the restoration of recombination even in situations in which reversion could be selected. When the number of potential breakpoints is relatively high, the probability of partially overlapping inversions occurring is higher than the probability of a reversion occurring, regardless of the relative rates of inversions and deleterious mutations.

The reversion of inversions has been reported only in very rare cases in which inversions occur at specific breakpoints rich in repeated elements (*51, 52*), as in our simulations in which only a few possible breakpoints were present in the genome. The number of genomic positions at which inversions can occur in natural conditions is unknown, but this number is likely to be high given the chaos of rearrangements observed in some sex and mating-type chromosomes with hundreds of different breakpoints (*16, 45, 53, 54*). A recent study reported the recurrent appearance of inversions at the same positions, but also the existence of numerous potential breakpoints of inversions in human genomes (*45*). Moreover, chromosomal rearrangements rapidly accumulate in recently established regions of non-recombination (*53*) and overlapping inversions are observed on many sex chromosomes, which should prevent the restoration of recombination by reversions (*16, 55–58*). Therefore, over a wide range of realistic parameter values, the reversion of inversions should not prevent the stepwise formation of non-recombining sex chromosomes. When there are few potential inversion breakpoints in the genome, the mechanism stabilizing inversions through dosage compensation considered in a recent model (*46*) may act in addition to the mechanism of sex-chromosome evolution studied here, as these two mechanisms are not mutually exclusive. However, by contrast to the framework of our model, dosage compensation mechanisms require XY-like asymmetry and cannot, therefore, explain evolutionary strata on fungal mating-type chromosomes or bryophyte UV sex chromosomes (*5, 54*).

The theory proposed here applies to any locus with at least one permanently or nearly permanently heterozygous allele. It requires only that, within a population, individuals carry different numbers of partially recessive deleterious mutations in a genomic region that can be subjected to recombination suppression. This situation is probably frequent in diploid, dikaryotic or heterokaryotic organisms. For instance, the mean distance between two heterozygous positions is ~1000 bp in primates (*59*), and it has been estimated that most of mutations are probably partially recessive and deleterious (*22, 60*). Chromosomal rearrangements spanning hundreds of kilobases are frequently observed (*45, 61, 62*), and would, therefore, be expected to capture several deleterious variants.

We show that inversions are more likely to spread in regions harboring many segregating deleterious mutations, in which they have a greater chance of capturing highly advantageous haplotypes. Our model therefore predicts that species harboring a large number of deleterious recessive variants, due to their mating system or high mutation rate, for example, will be more prone to the evolution of large non-recombining regions with evolutionary strata on sex chromosomes than species with a low mutation load. Furthermore, in species with large population sizes, the time required for an inversion to become fixed may exceed the time for which the number of deleterious mutations in the inversion remains below average. This would prevent some inversions from becoming fixed, potentially lowering the rate of expansion of non-recombining regions on sex chromosomes in species with large population sizes. Variations in population size, mutation rates and mating systems across lineages may, therefore, account for the large number of different sex-chromosome structures in nature, with some organisms maintaining homomorphic sex chromosomes and others evolving highly differentiated sex chromosomes with multiple strata (*1*). In plants, for example, selection at the haploid stage seems to result in the efficient purging of recessive deleterious mutations (*63*), potentially accounting for the smaller non-recombining region observed on the sex chromosomes of plants and algae than in animals (*64, 65*). In fungi, evolutionary strata around mating-type genes have been reported only in species with an extended dikaryotic stage (*5*). This is consistent with our model, in which recombination suppression evolves through the sheltering of deleterious mutations, which cannot occur in haploid organisms.

In conclusion, given its simplicity and wide scope of application, our model of sex chromosome evolution is a powerful alternative to other explanations (*6*), although the various theories are not mutually exclusive. The strength of our model lies in the absence of strong assumptions, such as sexually antagonistic selection, XY asymmetry or small population sizes. Our model, based on the often overlooked observation that recessive deleterious mutations are widespread in genomes within natural populations, can explain the evolution of stepwise recombination suppression over a wide range of realistic parameter values. Furthermore, it can also explain why some supergenes, such as those in butterflies and ants, display evolutionary strata (*13, 14*), and why many fungal mating-type loci display a stepwise extension of non-recombining regions despite the absence of sexually antagonistic selection or XY-like asymmetry in these organisms (*5, 16*). Our model therefore provides a general and simple framework for understanding the evolution of non-recombining regions around loci carrying permanently heterozygous alleles.

## Materials and Methods

### Infinite population deterministic model

We consider the discrete-time evolution of an infinite size, randomly mating population experiencing only deleterious recessive mutations, with heterozygotes and homozygotes suffering from a *1-hs* and *1-s* reduction in fitness, respectively. At all *n* sites, mutations are at the same mutation-selection equilibrium frequency, denoted *q*. We used non-approximated values of *q* as derived by (*66*):

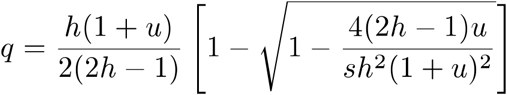

We assume that the sites are independent. The number of mutations carried by a chromosomal segment of length n then follows a binomial distribution of parameters (*n, q*). We follow the frequency of an inversion I of size *n*, considering that it captures *m* mutations and that it appears in a population carrying only non-inverted segments. The mean fitness of a non-inverted homozygote can be computed as follows (see Appendix, section 2):

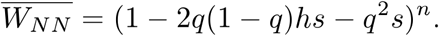

Note that in the parameter regimes where we can make the approximation *q* ≈ *u/hs* with *q* << 1, we have 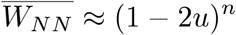, in accordance with mutation load theory (*34, 67*). Similarly, an individual who is heterozygous for an inversion with *m* mutations has a mean fitness of:

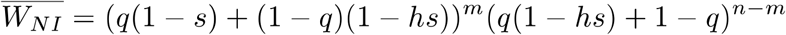

Assuming that *q* << 1, Nei et al., 1967 (*28*), considered that individuals heterozygous for an inversion had no mutations homozygous, such as 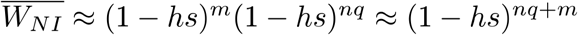. Note that we use non-approximated values in computations. An individual who is homozygous for a segment I with *m* mutations is homozygous for all these mutations. Its fitness can therefore be expressed as:

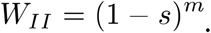

The inversion frequency trajectory can be determined with a simple two-locus two-allele model. We considered four different situations, depending on the possible heterozygosity at the locus with permanently heterozygous alleles. The Appendix, sections 4-8, presents the evolution in time of the frequency of the inversion in detail in these four situations. Here, we briefly describe the results for inversions more or less linked to a permanently heterozygous allele in a XY system. The change in frequency of inversions on the Y chromosome, on the X chromosome or on autosomes between generations *t* and *t* + 1 (Figure 2c) is described by

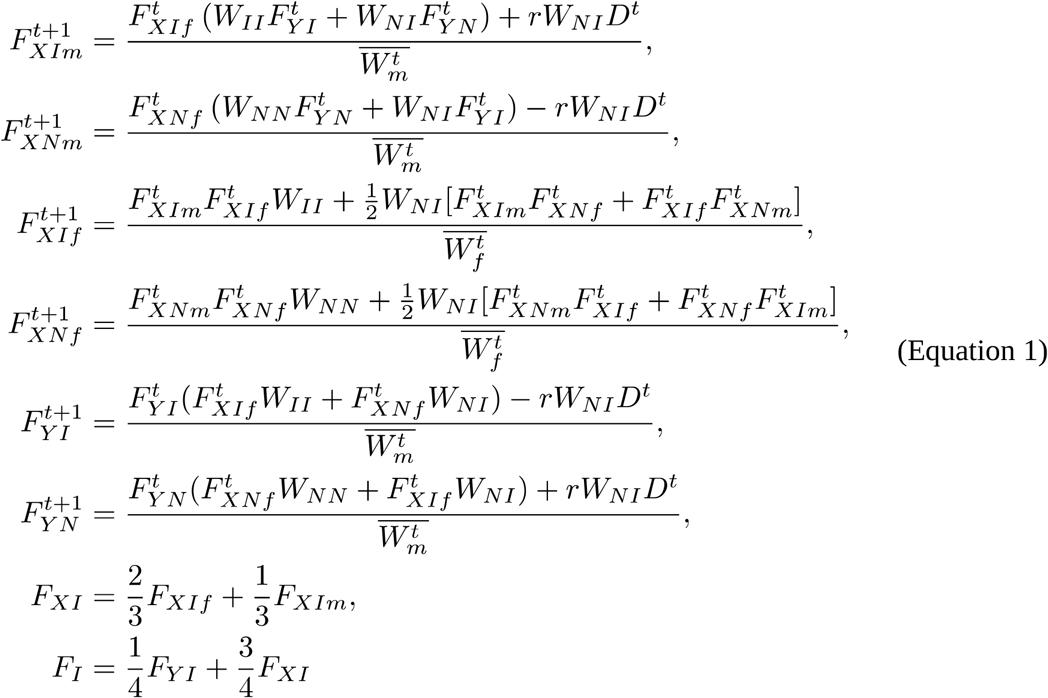

where 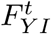 is the frequency of inversions in the population of Y chromosomes at time *t*, 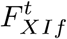 is the frequency of the inversion on the X chromosome in females (respectively 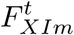 in males), *r* is the rate of recombination between the inversion and the sex-determining locus, *D* is their linkage disequilibrium and 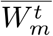 is the mean male fitness (Figure 2c; see Appendix, section 8 for details). When *r*=0.5, this system of equations describes the evolution of inversions on autosomes. Unless stated otherwise, the deterministic simulations presented here (Figures 2c and S4-8) were performed with an initial *D*=-0.01 or *D*=0.01, depending on the allele which the inversion appeared linked to. Results for various initial linkage disequilibrium values are presented in Figure S6. When the inversion captures the male-determining allele, *r*=0, *F_XIm_* = *F_XIf_* = 0, and *F_XNf_* = *F_XNm_* = 1 at any time *t*. The equations for 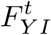, the frequency of inversions capturing the male-determining allele then reduces to

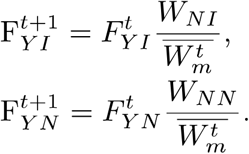

with 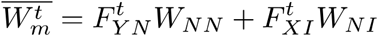. Substituting *F_YN_* by 1 − *F_YI_* and withdrawing 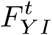, we have

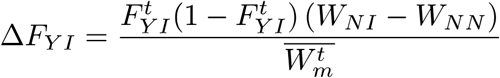

The search for *F_YI_* such that Δ*F_YI_* = 0 readily gives two equilibria : 0 and 1. Since 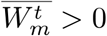 and 0 ≤ *F_YI_* ≤ 1, we can conclude that:

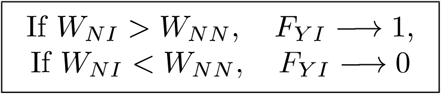

In the case of an inversion appearing on an autosome (*r*=0.5), *F_XIm_* = *F_XIf_* = *F_YI_* and 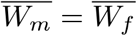, so that the change in inversion frequency in the population is:

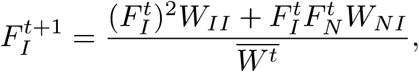

and

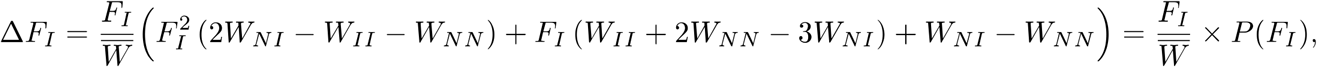

with *P*(*X*) = *X*^2^(2*W_NI_* − *W_II_* − *W_NN_*) + *X* (*W_II_* + 2*W_NN_* − 3*W_NI_*) + *W_NI_* − *W_NN_*.

The equilibria are 0 and the roots of the polynomial P, which are 1 and 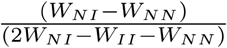. We thus have:

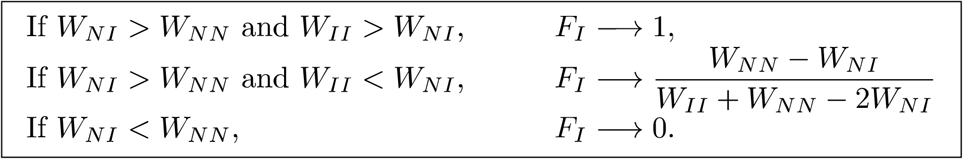

Inversion equilibrium frequencies therefore depend, as expected, on the relative fitness of homozygotes and heterozygotes for the inversion and of the non-inverted homozygotes. We thus derive the conditions for the inversion to be favoured or disfavoured as a function of the number of mutations captured by the inversion (*m*). A straightforward computation gives (see the Appendix, section 9):

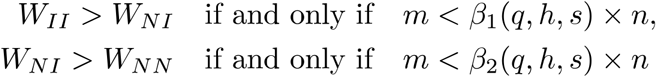

with

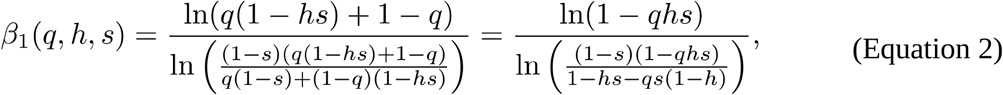

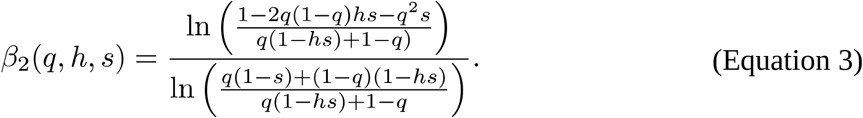

Assuming *q*<<1 and *s*<<1, these quantities can be approximated by 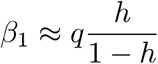 and *β*_2_ ≈ *q* [the latter in accordance with a previous study (*28*)]. When *qhn*/(1 − *h*) < *m* < *nq*, inversions should thus go to fixation on Y chromosomes and stabilize at intermediate frequency on autosomes. When *m* < *qhn*/(1 − *h*), inversions should go to fixation on autosomes and on the Y chromosome. Observe that the closer *h* is to ½ (i.e., the scenario without dominance), the smaller the difference between the thresholds *β*_1_ and *β*_2_ is. When h=0.5, *β*_1_ = *β*_2_, inversions should therefore go to fixation on autosome when they have fewer mutations than average, as they have then no homozygous disadvantage preventing their fixation. In contrast, when *h* is small, *β*_1_ is significantly smaller than *β*_2_, showing that the condition for heterozygotes to be favoured over non-inverted homozygotes (*W_NI_* > *W_NN_*) is much easier to meet than for inversion homozygotes to be favoured over inversion heterozygotes (*W_II_* > *W_NI_ W_II_* > *W_NI_*): inversions are therefore much likely to be maintained at intermediate frequencies on autosomes than to fix. See Figure S1 for a graphical representation of these results and the Appendix, section 9, for derivation details.

We used these equilibrium frequencies as functions of *m* to compute the expected equilibrium frequency of inversions that can occur in the genome (*F_equ_*, Figure 2d and S9). To do so, we sum, for all m ∈ {0, …, *n*} the equilibrium frequency of inversions (*F_equ,m_*) weighted by their occurrence probability (*P_m_*). For inversions on the Y chromosome, we therefore have:

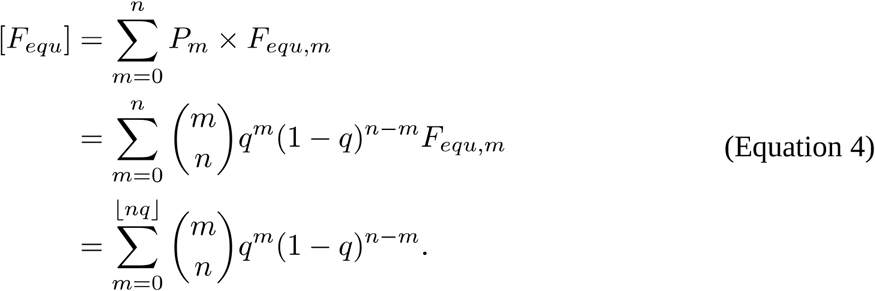

For inversions on autosomes, we obtain, using the approximate values for *β*_1_ and *β*_2_ derived earlier in the case q<<1 and s<<1:

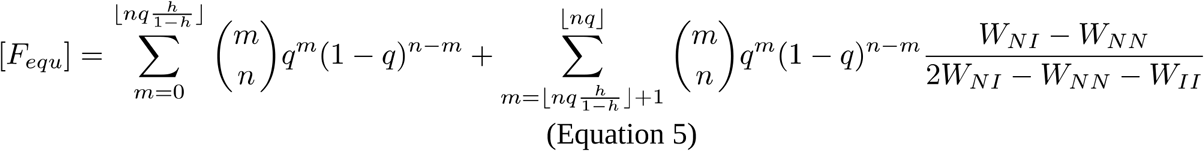

The expected equilibrium frequency of less-loaded inversions (Figure 2d) is therefore the expected equilibrium frequency of all inversions divided by the probability of occurrence of less-loaded inversions:

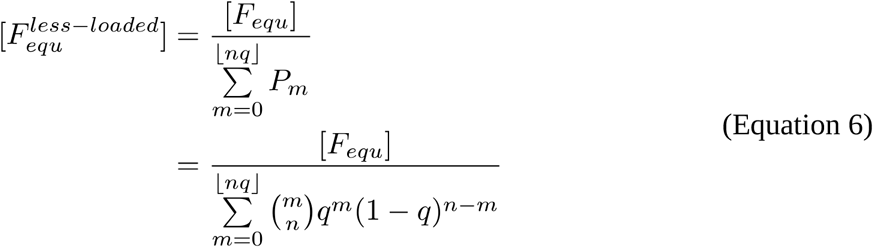

### Individual-based simulations

We used SliM V3.2 (*68*) to simulate the evolution of a single panmictic population of *N*=1000 or *N*=10,000 individuals in a Wright-Fisher model. To assess the fate of inversions under various conditions (Figure 3), we simulated individuals with two pairs of 10Mb chromosomes on which mutations occurred at a rate *u*, with *u* ranging from 10^-6^ to 10^-9^ per bp, their dominance coefficient *h* ranged from 0 to 0.5 (0, 0.01, 0.1, 0.2, 0.3, 0.4, 0.5) and their selection coefficient *s* from −0.5 to 0 (0, −0.001, −0.01, −0.1, −0.25, −0.5). The main and supplementary figures show the results for *u*=10^-8^ and *u*=10^-9^ unless otherwise stated. We also simulated populations where each occurring mutation had its selection coefficient *s* drawn from a gamma distribution with a shape of 0.2, and its dominance coefficient *h* randomly sampled among 0, 0.001, 0.01, 0.1, 0.25, 0.5 with uniform probabilities (Figure S17). We considered recombination rates of 10^-6^ and 10^-5^ per bp, which gave similar results. Results are presented for analyses in which the recombination rate was 10^-6^. Among the 5,000,000 sites on chromosome 1, a single segregating locus was subject to balancing selection, for which several situations were considered: (i) the locus had two alleles, only one of which was permanently heterozygous, mimicking a classical XY (or ZW) determining system (Figure 3), (ii) the locus had two permanently heterozygous alleles, mimicking, for instance, the situation encountered at most fungal mating-type loci, iii) the locus had three (or more) permanently heterozygous alleles, mimicking, for instance, the situation encountered in plant self-incompatibility systems and mushroom (*Agaromycotina*) mating-type loci.

For each parameter combination (*u, h, s, N*, heterozygosity rule at the locus with a permanently heterozygous allele), a simulation was run for 15,000 generations, to allow the population to reach an equilibrium for the number of segregating mutations. Figure S18 shows that each population had reached equilibrium by the end of the burn-in period. The population state was saved at the end of this initialization phase. These saved states (one for each parameter combination) were repeatedly used as initial states for studying the dynamics of recombination modifiers. Recombination modifiers mimicking inversions of 500kb, 1000kb, 2000kb and 5000kb were then introduced on chromosome 1 around the locus under balancing selection (X-linked or Y-linked inversions) or on chromosome 2 (autosomal inversions). For each parameter combination (*h, s, u, N*, heterozygosity rule, size of the region affected by the recombination modification and position on the genome), we ran 10,000 independent simulations starting with the introduction of a single recombination modifier in the same saved initial population. These inversion-mimicking, recombination modifier mutations were introduced on a single, randomly selected chromosome and, when heterozygous, they suppressed recombination across the region in which they reside (i.e., as a *cis*-recombination modifier). We monitored the frequency of these inversion-mimicking mutations during 10,000 generations, during which all evolutionary processes (such as point mutation, recombination and mating) remained unchanged, e.g. mutations were still appearing on inversions following their formation. Under the same assumptions and parameters, we also studied the dynamics of recombination modifiers suppressing recombination also when homozygous and not only when heterozygous, again across a fragment in which they reside (Figure S14).

To study more specifically the formation of evolutionary strata on sex chromosomes (Figures 4 and 5), we also simulated the evolution of two 100 Mb chromosomes, one of which carried an XY sex-determining locus at the 50 Mb position, over 115,000 generations (including an initial burn-in of 15,000 generations): individuals could be either XX or XY and could only mate with individuals of a different genotype at this locus. We simulated randomly mating populations of *N*=1000 and *N*=10,000 individuals. Point mutations appeared at a rate of *u*=10^-9^ per bp, and their individual selection coefficients were determined by sampling a gamma distribution with a mean of −0.03 and with a shape of 0.2; these parameter values were set according to observations in humans (*22, 69*). For each new mutation, a dominance coefficient was chosen from the following values, considered to have uniform probabilities: 0, 0.001, 0.01, 0.1, 0.25, 0.5. At the beginning of each simulation, we randomly sampled *k* genomic positions over the two chromosomes that could be used as inversion breakpoints, with *k* being 10, 100, 1000 or 10,000. After the 15,000 generations of the burn-in period allowing populations to reach an equilibrium in terms of the number of segregating mutations, we introduced each generation *j* inversions in the population, *j* being sampled from a Poisson distribution of parameter *λ*, with *λ* = *N* ∗ *k* ∗ *u_i_*, *u_i_* being the inversion rate. In order to keep the simulation time tractable, we used inversion rates allowing one inversion to occur on average each generation in the population (i.e. *N* ∗ *k* ∗ *u_i_* = 1). For each inversion, the first breakpoint was randomly chosen among the *k* positions, and the second among the potential breakpoint positions less than 20Mb apart on the same chromosome (considering therefore only a subset of the *k* positions for the second breakpoints). Two independent inversions could use the same breakpoints, allowing in particular inversion reversion restoring recombination. We assumed that inversions subsequently partially overlapped by another inversion or that captured a smaller inversion could not reverse, i.e. the restoration of recombination by further inversions, even if using the same breakpoints, was prevented. For *N*=1000, for each values of *k* (i.e. 10, 100, 1000, 10000), we ran 10 simulations (Figure 5). For *N*=10,000, because of computing limitation (each simulation taking two or three weeks to run), we only ran one simulation per number of breakpoints.

Simulations were parallelized with GNU Parallel (*70*).

## Acknowledgment

We thank Ricardo Rodriguez de la Vega, Fanny Hartmann, Jacqui Shykoff, Sylvain Billiard, Janis Antonovics and Olivier Tenaillon for comments on a draft version of the manuscript. ET thanks Denis Roze for insightful discussions. ET and AV acknowledge support from the chaire program « Mathematical modelling and biodiversity » (Ecole Polytechnique, Museum National d’Histoire Naturelle, Veolia Environnement, Fondation X). PJ thanks the authors of SLiM for their outstanding software and manual.

## Author contributions

Original ideas, PJ and TG; deterministic model conception, PJ and ET with input from AV; simulations and data analyses, PJ; interpretation, PJ, TG and ET; manuscript writing, PJ and TG; editing, ET and AV; project management and funding, TG.

## Data and materials availability

The SLiM and R scripts used for this study are available from GitHub (https://github.com/PaulYannJay/SexChromosomeTheory). No new data were generated for this study. Details concerning the mathematical modeling are available in the appendix.

## List of supplementary materials

Figures S1-19

Appendix (supplementary methods)

