## Supplementary figures for "Modeling the stepwise extension of recombination suppression on sex chromosomes and other supergenes through deleterious mutation sheltering"

**This document contains:**
Figures S1-19

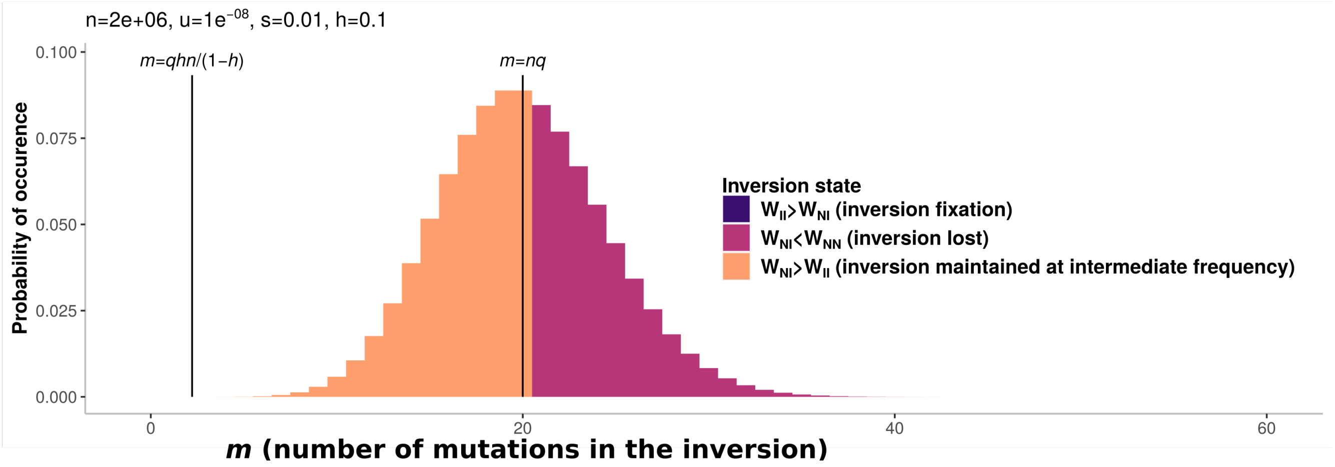


**Figure S1 | Distribution of the occurrence probability of inversions depending on the number of mutations they capture.**

This figure represents the binomial distribution of the number of mutations on a segment of *n*=2,000,000 sites, with mutations occurring at a rate *u*=1e-08 and having selective effect of *s*=0.01 and *h*=0.1. The filling color of the distribution represents the expected state of the inversions on autosomes based on equations 2 and 3 (Methods). The probability of occurrence of an inversion carrying fewer than $m=qhn/(1-h)$ mutations, and therefore fixing on the autosome (dark blue filling), is so low that it is not visible on this plot.

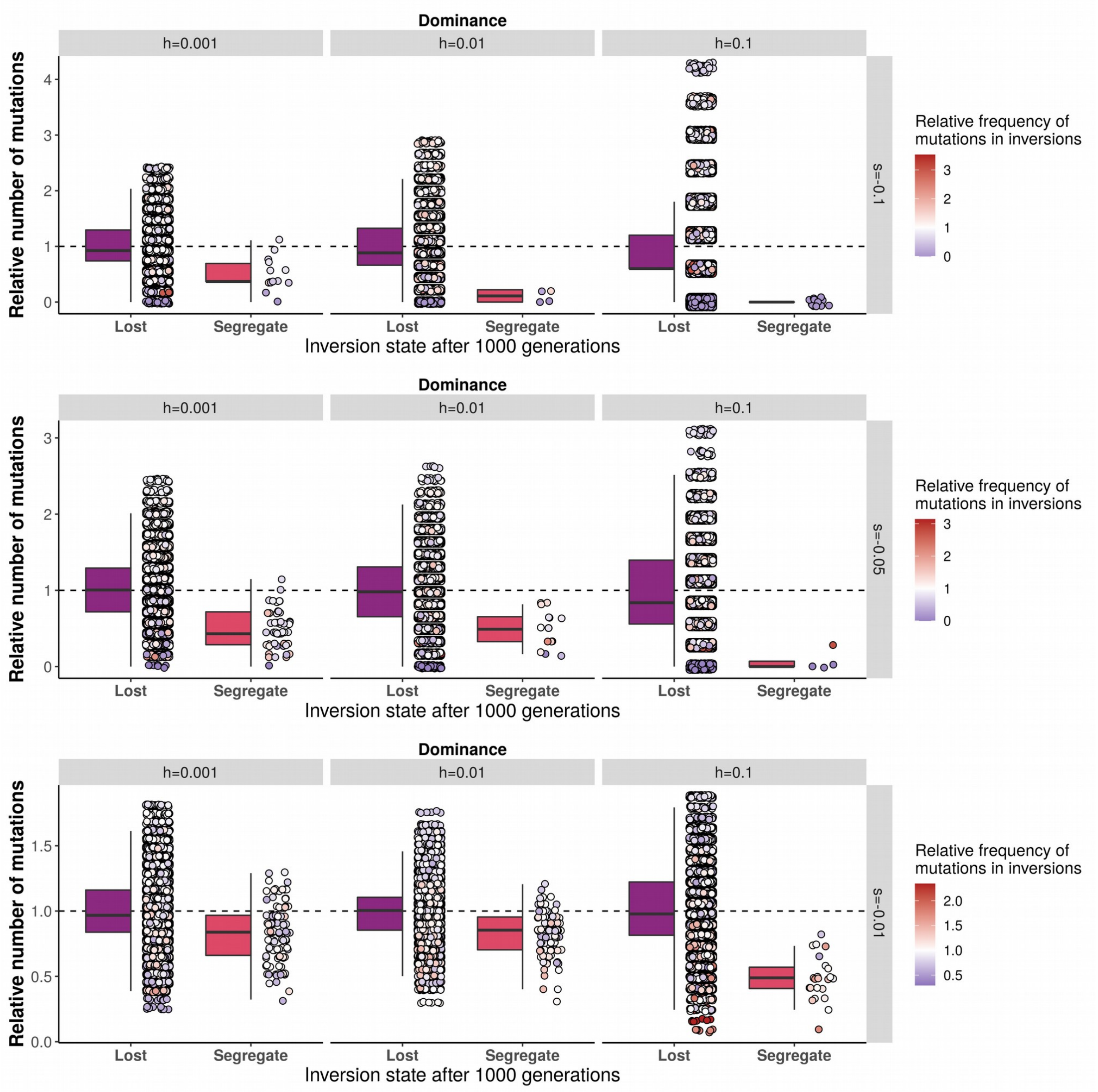


**Figure S2. Fate of neutral chromosomal inversions on autosomes depending on the relative number of mutations they carry compared to population average and the relative frequency of these mutations**
Each dot represents a 2Mb inversion on an autosome. For each parameter combination, the fate (i.e., being lost or still segregating) of 10,000 inversions after 1000 generations in stochastic simulations of a population of N=1000 individuals is displayed depending on the number of mutations they captured upon formation relative to the population average. Only inversions not linked to a permanently heterozygous allele were considered. Results for sex-linked inversions are displayed in Figure S3. The Y axis represents the number of mutations captured by the inversions upon formation relative to the mean number of mutations within the same region in non-inverted segments. Dot color represents the mean of the frequencies (in the entire population) of the mutations captured by inversions relative to the mean of the frequencies (in the entire population) of mutations in non-inverted segments: a blue dot represents an inversion capturing mutations rarer than average whereas a red dot represents an inversion capturing mutations more frequent than average. Boxplot elements: central line: median, box limits: 25th and 75th percentiles, whiskers: 1.5x interquartile range. The vast majority of inversions still segregating after 1000 generations carried fewer or rarer mutations than the population average.

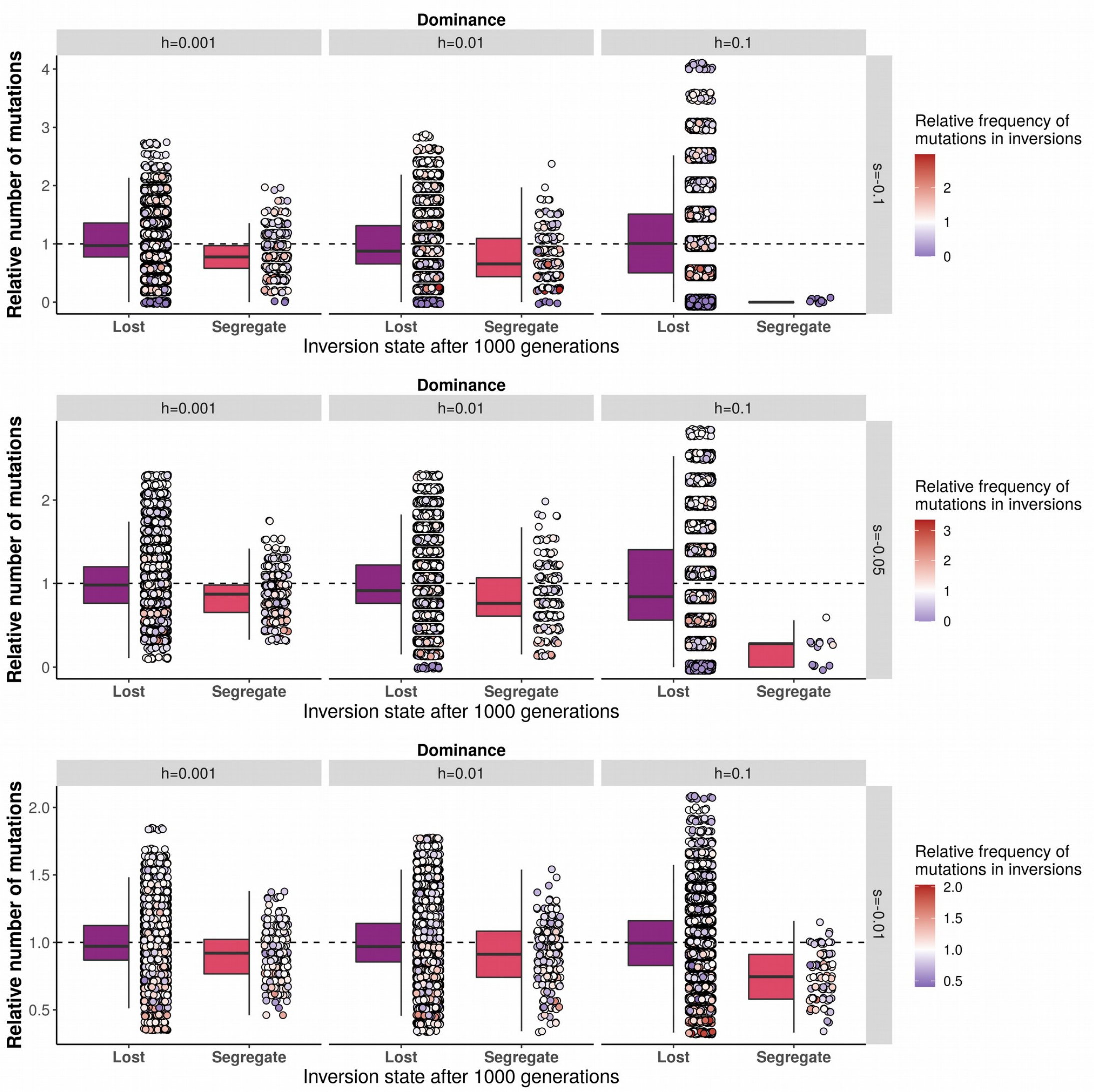


**Figure S3. Fate of neutral chromosomal inversions linked to a permanently heterozygous allele depending on the relative number of mutations they carry compared to population average and the relative frequency of these mutations**

Similar to Figure S2 but with inversions linked to a permanently heterozygous allele (such as a Y sex-determining allele). For each parameter combination, the fate (i.e., being lost or still segregating) of 10,000 inversions after 1000 generations in stochastic simulations is displayed depending on the number of mutations captured upon formation relative to the population average. Dot color represents the mean of the frequencies (in the entire population) of the mutations captured by inversions relative to the mean of the frequencies (in the entire population) of mutations in non-inverted segments: a blue dot represents an inversion capturing mutations rarer than average whereas a red dot represents an inversion capturing mutations more frequent than average. Boxplot elements: central line: median, box limits: 25$^{th}$ and 75$^{th}$ percentiles, whiskers: 1.5x interquartile range. The vast majority of inversions still segregating after 1000 generations carried fewer or rarer mutations than the population average.

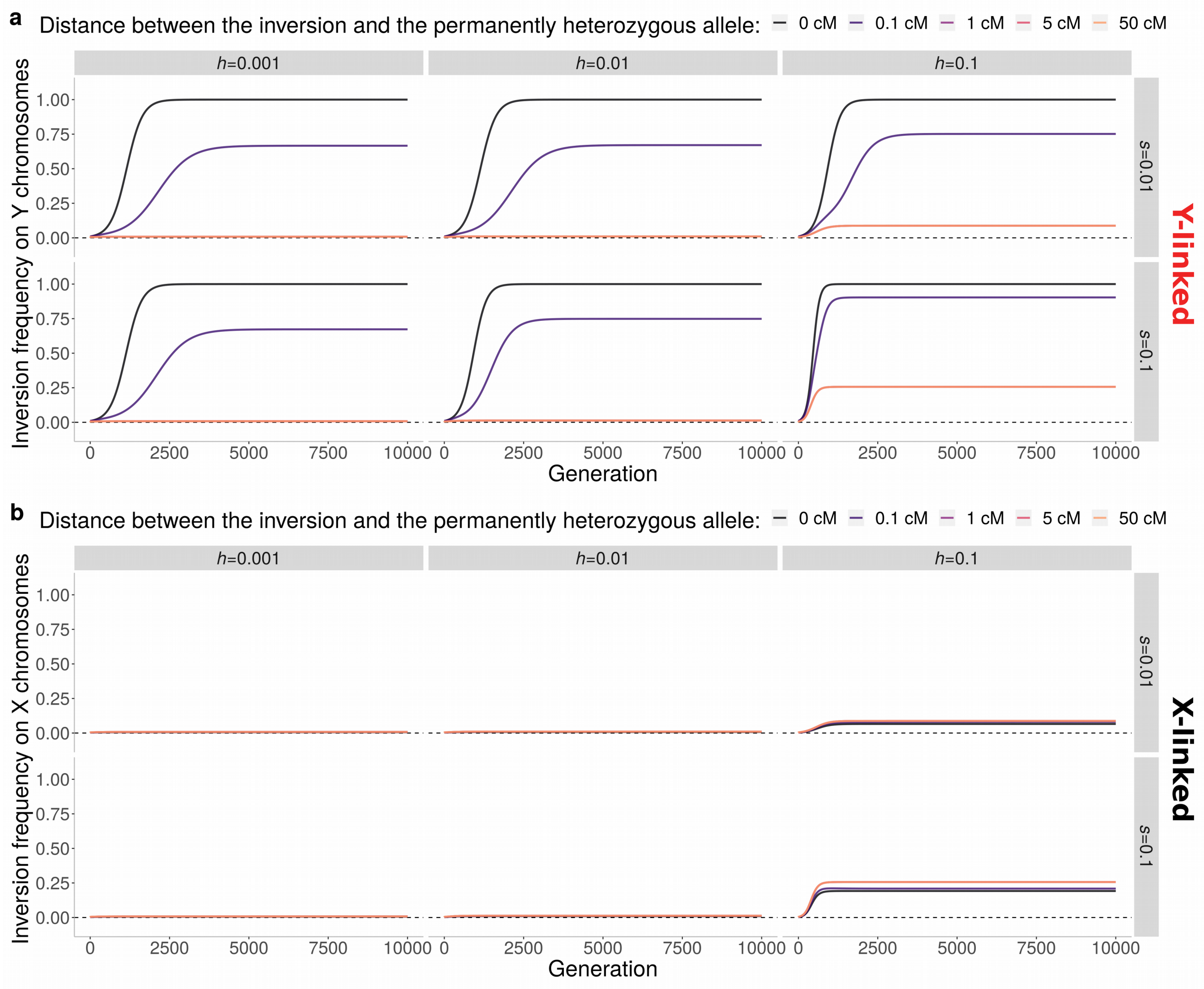


**Figure S4. Deterministic trajectories of inversion frequency when in linkage with the sex-determining allele on the Y chromosome or on the X chromosome.**

Similar to Figure 2c, but considering both X-linked and Y-linked inversions, and various recombination rates between the sex-determining locus and inversions. Trajectory of inversions are represented for different values of the selection and dominance coefficients of their mutations and the linkage of these inversions to a X or to a Y sex-determining allele. Mutation frequencies are at mutation-selection equilibrium (see methods) with a mutation rate of $u=10^{-8}$. Inversions of 2Mb are introduced with an initial linkage disequilibrium of $D=0.01$ or $D=-0.01$, depending on whether they appear linked to the male-determining allele on the Y chromosome or to the female-determining allele on the X chromosome. The figure illustrates the case of inversions carrying a number of mutations 20% lower than the population average ($m=[0.8*nq]$), as frequently observed (Figures S1-3). **a,** Inversions appearing in linkage with a sex-determining allele on the Y chromosome (permanently heterozygous). The frequency of the inversion in the population of Y chromosomes is followed during the first 10000 generations. Linkage strength is indicated by line colors (distance in cM between inversions and the sex-determining allele). Inversions at 50cM from the sex-determining locus behave like inversions on autosomes. There is a nearly perfect overlap between curves representing the 1, 5, and 50 cM linkage cases, so that only 50 cM curves are visible. **b,** Inversions appearing in linkage with the sex-determining allele on the X chromosome. The frequency of the inversion in the population of X chromosomes is followed during the first 10000 generations. Linkage strength is indicated by line colors (distance in cM between inversions and the sex-determining allele). See Appendix, section 8, for modelling details.

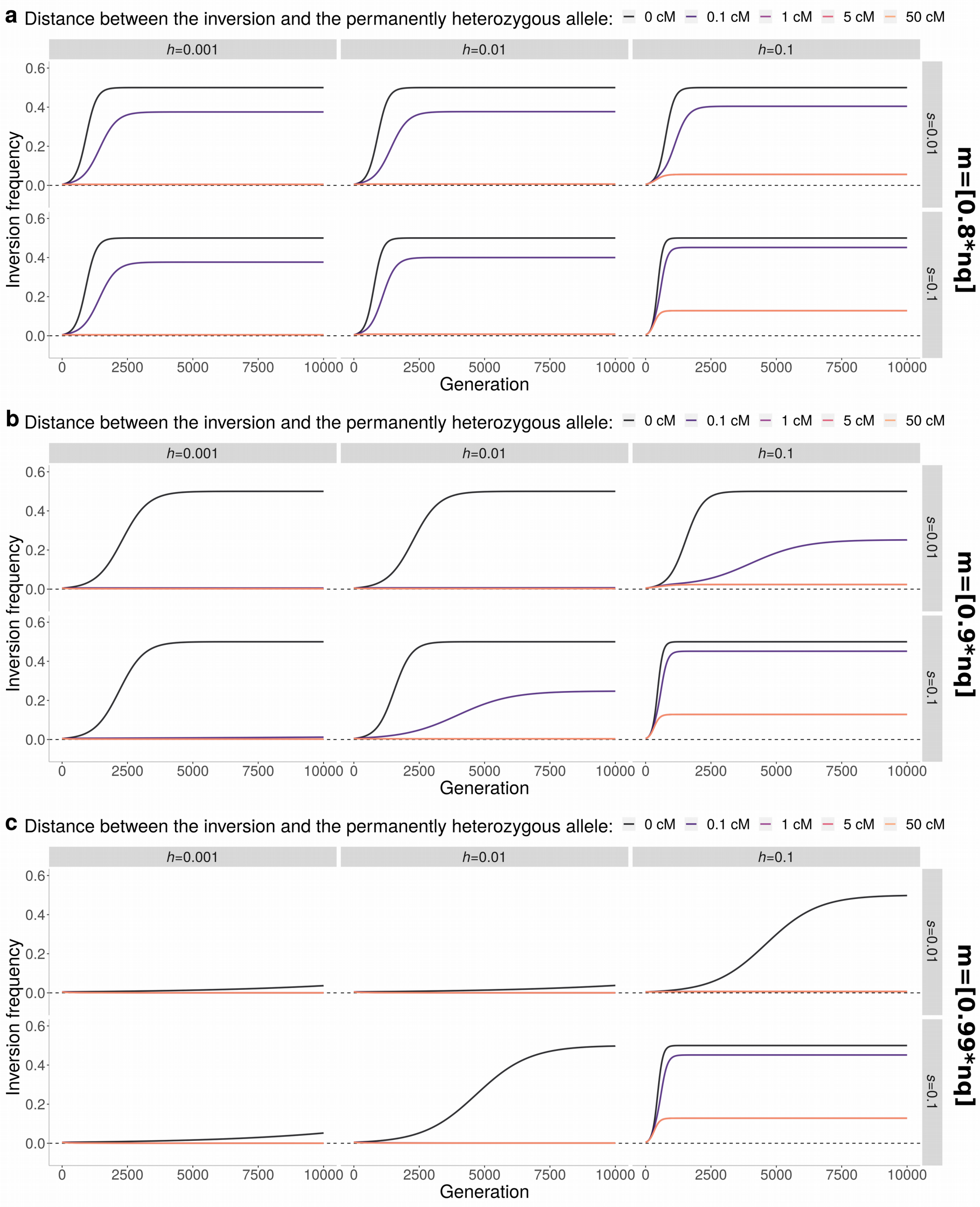


**Figure S5. Deterministic trajectory of inversion frequency depending on their initial mutation load and their linkage to a locus with two permanently heterozygous alleles.**

Similar to Figures 2c and S4, but considering inversions in linkage to a locus with two permanently heterozygous alleles (and not only one as in the XY systems), and various mutation loads associated with the inversions upon formation. Trajectory of inversions in an infinite population depending on the selection coefficient of the segregating mutations, the dominance coefficient of these mutations and the linkage of these inversions to a permanently heterozygous locus with two alleles. Mutations are at mutation-selection equilibrium frequencies (see Methods) with a mutation rate of $u$=$10^{-8}$. Inversions of 2Mb are introduced with an initial linkage disequilibrium of $D$=0.01. Inversions are linked to only one of the two alleles at the permanently heterozygous locus, and so even fully linked inversions can increase up to a maximum frequency of 50%. Linkage strength is indicated by line colors (distance in cM between inversions and permanently heterozygous locus). Inversions at distances larger than 50cM from the sex-determining locus behave like inversions on autosomes. The figure illustrates the case of inversions carrying varying numbers of mutations compared to the population average. **a,** Case of inversions with 20% fewer mutations than the population average ($m=[0.80*nq]$). **b,** Case of inversions with 10% fewer mutations than the population average (m=[0.90*nq]). **c,** Case of inversions with 1% fewer mutations than the population average (m=[0.99*nq]). In all cases, there is a nearly perfect overlap between curves representing the 1, 5, and 50 cM linkage cases, so that only 50cM curves are visible. Individual-based simulations of the same system are shown in figure S14. See Appendix, section 6, for modelling details.

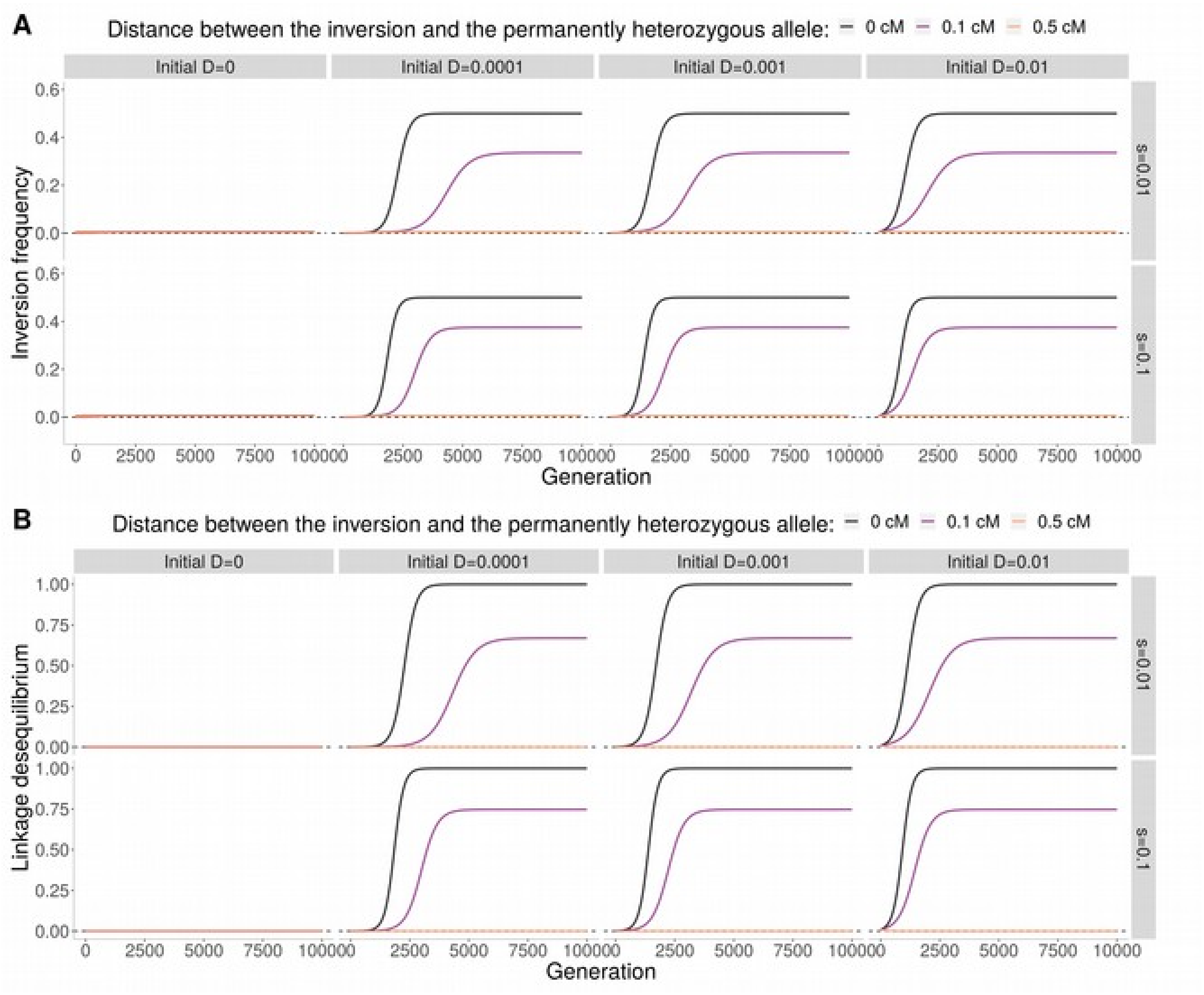


**Figure S6. Deterministic trajectory of inversion frequency depending on their linkage to a permanently heterozygous locus with two alleles (in centiMorgan) and on their initial association (initial linkage disequilibrium).**

Trajectory of inversions in infinite populations as a function of the selection coefficient of the segregating mutations, their linkage to a permanently heterozygous locus with two alleles, and the initial level of linkage disequilibrium (D) between the inversion locus and the permanently heterozygous locus. Mutations are at mutation-selection equilibrium frequencies. Inversions of 2Mb are introduced at different frequencies in the population, and with different levels of association with one of the two permanently heterozygous alleles, which therefore generates various initial linkage disequilibrium values (D). Inversions appear more or less linked to a locus with two permanently heterozygous alleles; completely linked inversions can therefore only increase up to a maximum frequency of 50%. Linkage strength is indicated by line colors, representing the distance in cM between inversions and the permanently heterozygous alleles/locus. Inversions at more than 50cM from the permanently heterozygous locus behave like inversions on autosomes. The figure illustrates the case of inversions carrying a number of mutations 20% lower than the population average ($m$=[0.8*$nq$]), as frequently observed (Figures S1-3). A dominance coefficient of $h$=0.01 was used for these deterministic simulations. **A,**

Inversion change in frequency. Similar to Figures 2c and S5 but considering inversions in linkage with a locus with two permanently heterozygous alleles, and various initial levels of linkage disequilibrium between the inversion locus and the permanently heterozygous locus. **B,** Change in the level of linkage disequilibrium ($D$) between the inversion locus and the permanently heterozygous locus. If there is no initial linkage disequilibrium (i.e., inversions are introduced at the same frequency in linkage with each of the two permanently heterozygous alleles), the inversion does not spread. As soon as there is an initial linkage disequilibrium, even very small, this linkage disequilibrium increases and the inversion spreads. Here, the initial linkage disequilibrium is artificially generated by introducing the inversions with different levels of association with a permanently heterozygous allele. In finite populations, this non-null initial linkage disequilibrium can easily be generated by founder-like effects (an inversion typically appearing on a single haplotype) or by the random fluctuations of inversion frequency induced by drift. See Appendix, section 6, for modelling details.

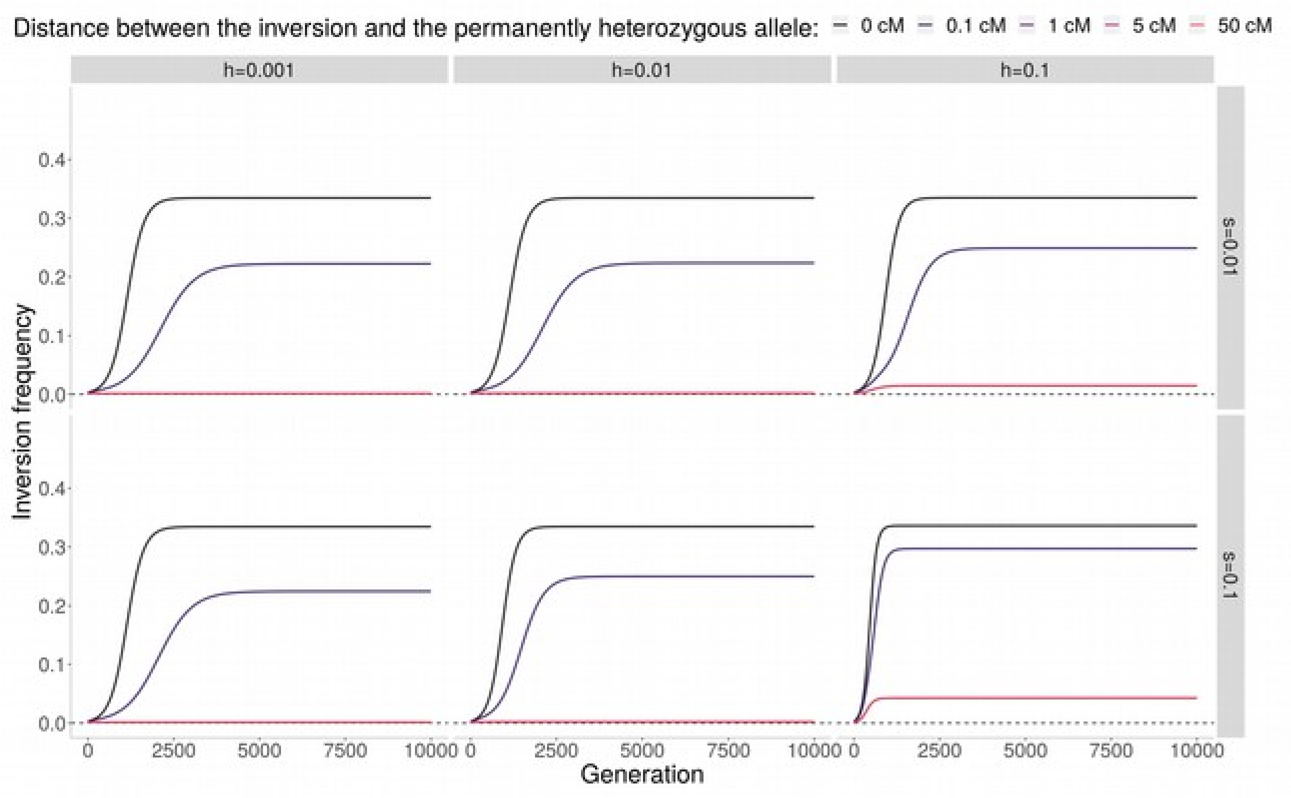


**Figure S7. Deterministic trajectory of inversion frequency when in linkage to a permanently heterozygous locus with three alleles.**

Similar to figure 2c and S3-5, but considering inversions in linkage to a locus with three permanently heterozygous alleles. Trajectory of inversions depending on the selection and dominance coefficients of their mutations and their degree of linkage to the permanently heterozygous locus. Mutation frequencies are at mutation-selection equilibrium (see Methods) with a mutation rate of $u$=$10^{-8}$. Inversions of 2Mb are introduced at a 0.01 frequency in linkage with one of the three permanently heterozygous alleles. The figure illustrates the case of inversions carrying a number of mutations 20% lower than the population average ($m$=[0.8*$nq$]), as frequently observed in simulations (Figure S1-3). Linkage strength is indicated by line colors, representing the distance in cM between inversions and permanently heterozygous alleles. When fully linked to the permanently heterozygous locus (0 cM), inversions can increase up to a maximal frequency of 1/3, meaning that they are fully associated to one of the three permanently heterozygous alleles. See Appendix, section 7, for modelling details.

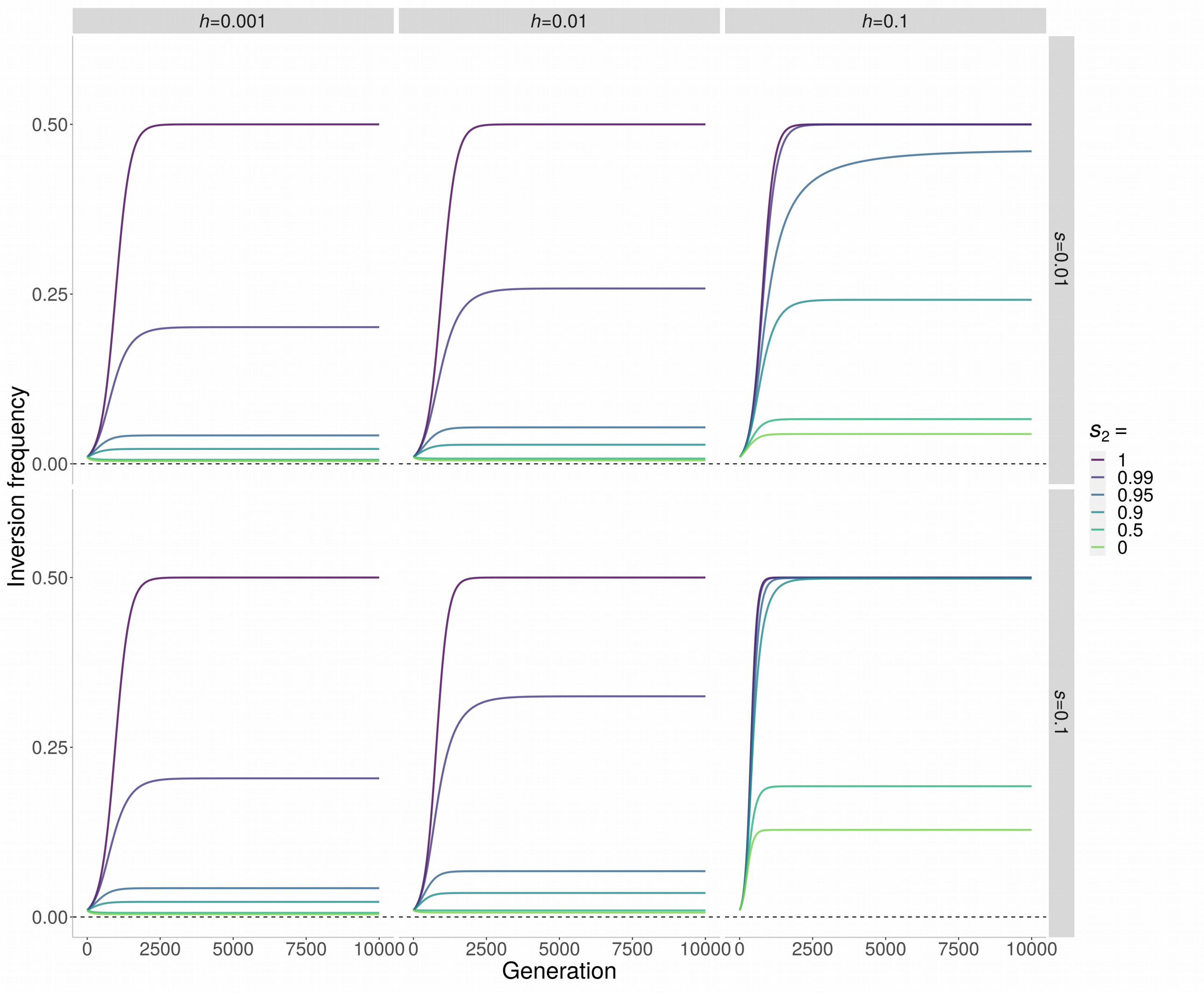


**Figure S8. Deterministic trajectory of inversion frequency when in linkage with an overdominant locus with no permanently heterozygous alleles.**

Similar to Figure 2c but with an overdominant locus with two alleles. Trajectory of inversions depending on the selection and dominance coefficients of their mutations and the strength of selection acting on the overdominant locus (with no permanently heterozygous alleles). Mutation frequencies are at mutation-selection equilibrium (see Methods) with a mutation rate of $u$=$10^{-8}$. The figure illustrates the case of 2Mb inversions carrying a number of mutations 20% lower than the population average ($m$=[0.8*$nq$]). Inversions appear in full linkage ($r$=0.0) with one the two alleles at an overdominant locus where both homozygotes have reduced fitness (1-$s_2$; when $s_2$ is close to 1, homozygotes suffer from a strong reduction in fitness and are therefore rare; See Appendix, section 5, for modelling details.).

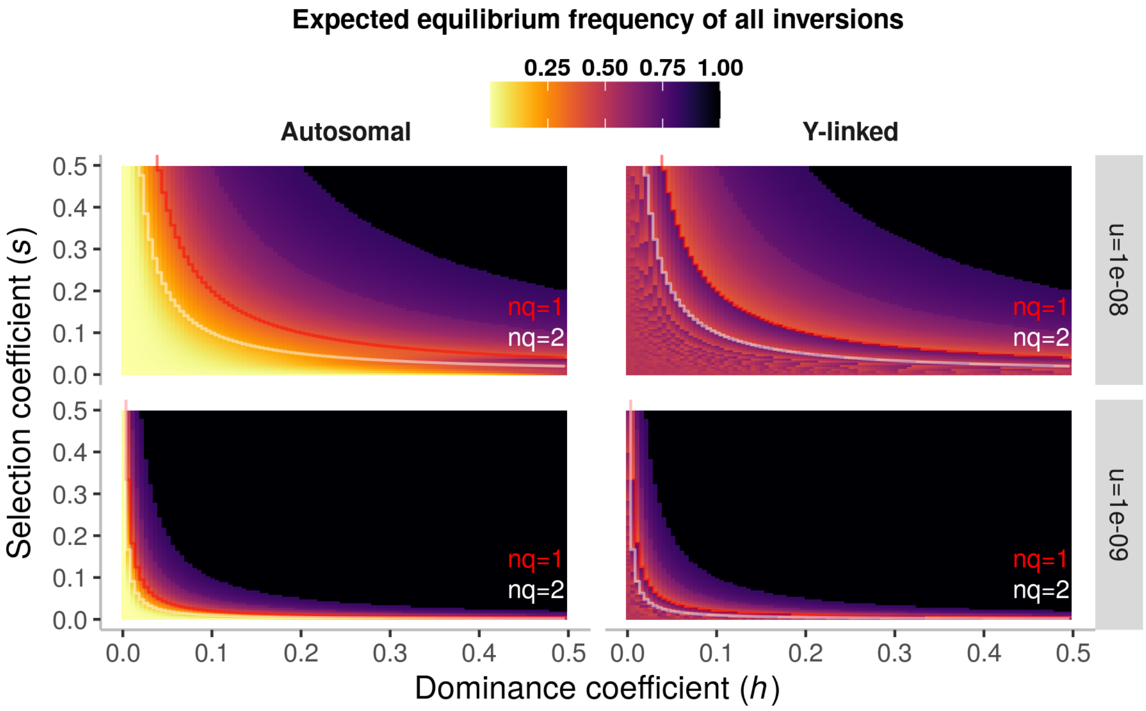


**Figure S9 | Expected equilbrium frequency of 2 Mb inversions as a function of selection and dominance coefficients.**

Expected equilibrium frequency of all 2 Mb inversions that can occur capturing the male-determining allele in an XY system (right column) or on an autosome (left column). Therefore, this is similar to figure 2d but considering all inversions, i.e. inversions less-loaded and more-loaded than average. The expected equilibrium frequency was calculated using equations 4 and 5 (Methods). The frequency displayed for Y-linked inversions is the frequency of inversions in the population of Y chromosomes. Y-linked inversions become fixed (equilibrium frequency=1) when $m<nq$ and are lost (equilibrium frequency=0) when $m>nq$ (see methods and main text). These two events have nearly equal probabilities when $nq$ is high (figure 2b and S1), and therefore the expected equilibrium frequency of Y-linked inversion is often around 0.5.

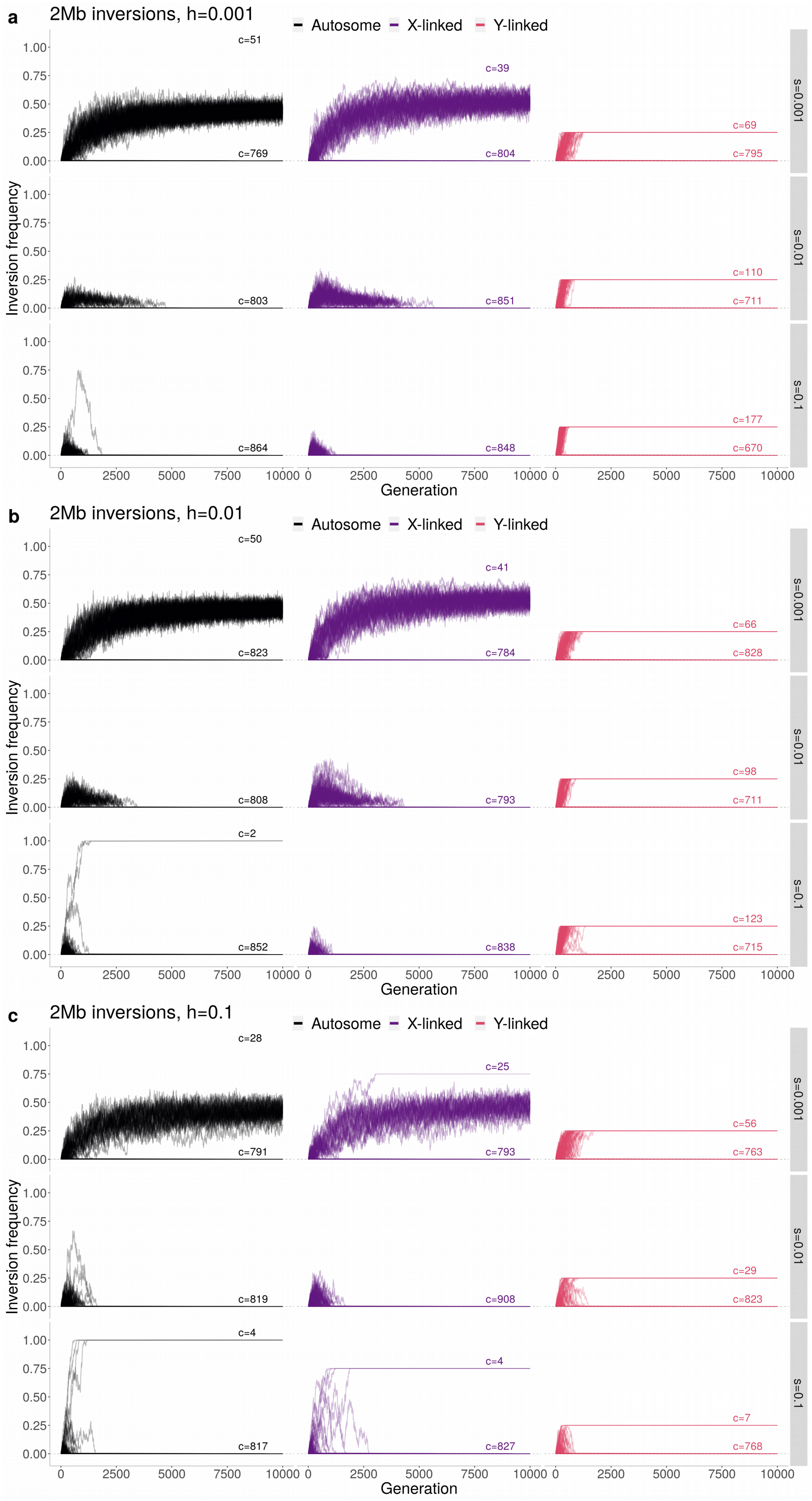
a
2Mb inversions, h=0.001
b
2Mb inversions, h=0.01
c
2Mb inversions, h=0.1
Autosome
X-linked
Y-linked
Inversion frequency
Generation
s=0.001
s=0.01
s=0.1
c=51
c=769
c=39
c=804
c=69
c=795
c=803
c=851
c=110
c=711
c=864
c=848
c=177
c=670
c=50
c=823
c=41
c=784
c=66
c=828
c=808
c=793
c=98
c=711
c=2
c=852
c=838
c=123
c=715
c=28
c=791
c=25
c=793
c=56
c=763
c=819
c=908
c=29
c=823
c=4
c=817
c=4
c=827
c=7
c=768

**Figure S10 | Evolution of 2Mb inversions in stochastic simulations during 10,000 generations.** Similar to Figure 3a but with other parameter values (*s*, *h*). For each parameter combination, 10,000 inversions of 2000kb were simulated in a population harboring an XY locus (two alleles, one being permanently heterozygous). Populations of 1000 individuals were simulated. We considered inversions fully linked to the Y sex-determining allele, fully linked to the X sex-determining allele or on an autosome. Only inversions not lost after 20 generations are displayed. In contrast to Figure 3a, the overall frequency of Y-linked and X-linked inversion is displayed (instead of the frequency of inversion in the Y chromosome population). Inversions fixed on the Y chromosome have therefore a 0.25 frequency (and 0.75 for X-linked inversions). For each parameter combination, the number of inversions lost and segregating or fixed at the end of the simulation is indicated above lines (« c=... »). Because N=1000, mutations with *s*=0.001 (i.e. 1/N) are nearly neutral, which explains that X-linked or autosomal inversions can spread in the population, in contrast to what is observed with *s*>0.001.

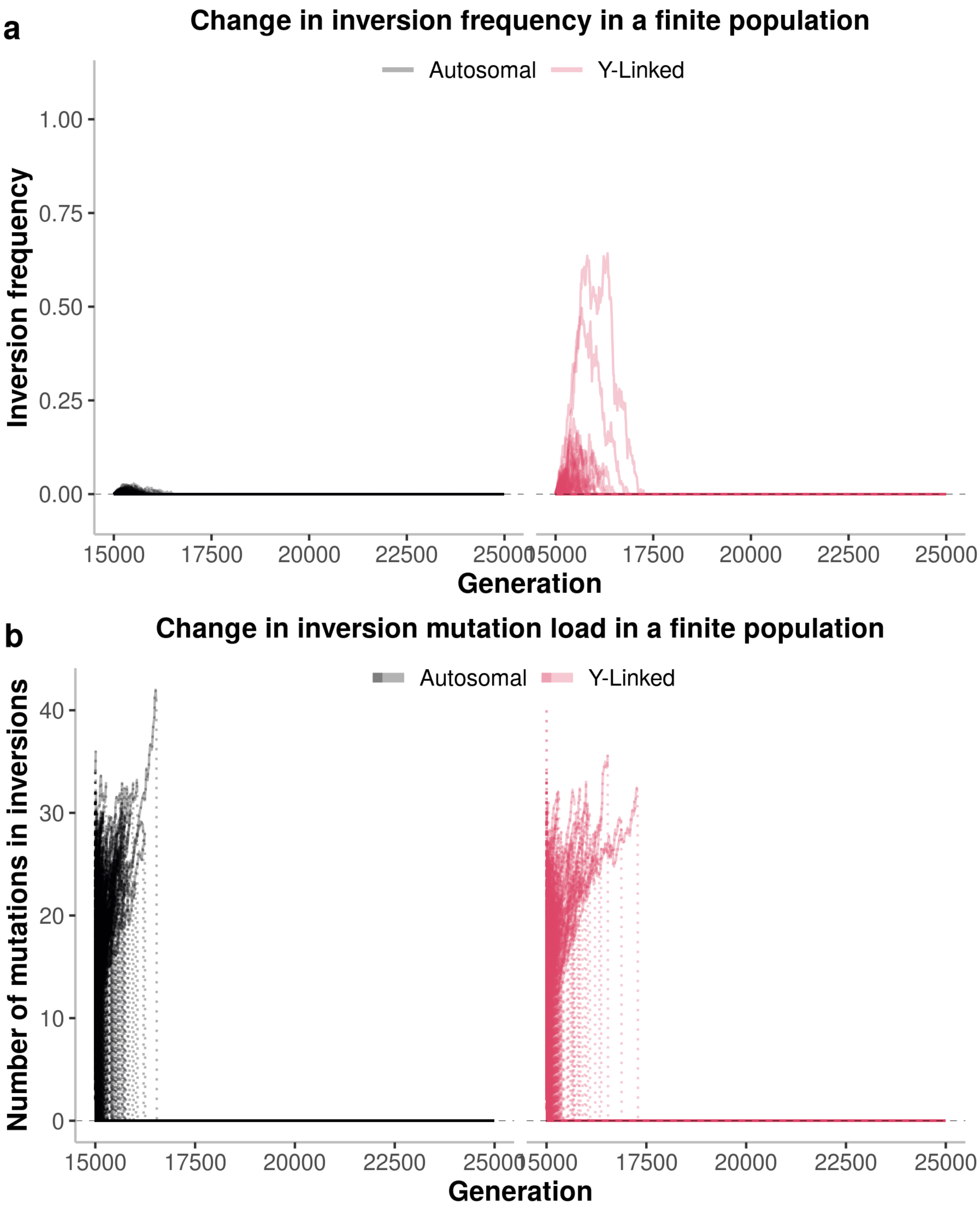


**Figure S11 | Evolution of 2Mb inversions in stochastic simulations over 10,000 generations in populations of *N*=10,000 individuals with *s*=0.01.**

Similar to Figure 3a-b but with *N*=10,000. **a,** Change in inversion frequency in stochastic simulations with 10,000 individuals experiencing recessive deleterious mutations at a rate *u*=1e-08, including following inversion occurrence, all mutations having the same dominance and selection coefficients (here, *h*=0.1 and *s*=0.01). The figure displays the frequency of inversions on an autosome and on a proto-Y chromosome, each line representing a specific inversion. **b,** Fluctuations in the mean number of mutations carried by the inverted segments in each of the 10,000 simulations, each line representing one simulation. In contrast to what happens with N=1000 (Figure 3a-b)**, s**ome inversions rise in frequency on the Y chromosome; however, the high population size renders the time for inversion fixation too long, they accumulate deleterious mutations before they can fix and therefore all inversions were lost at the simulation end.

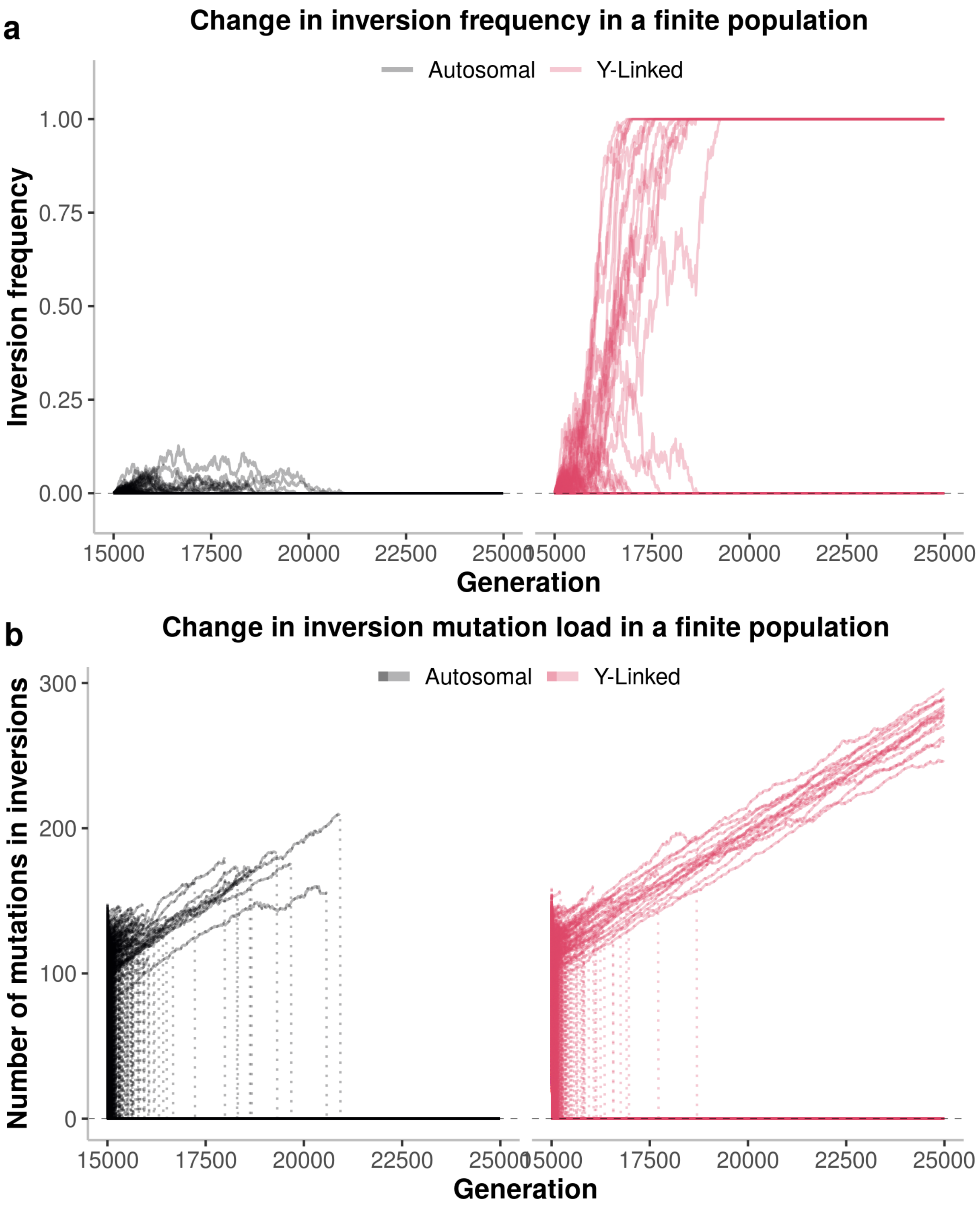


**Figure S12 | Evolution of 2Mb inversions in stochastic simulations over 10,000 generations in populations of *N*=10,000 individuals with *s*=0.001.**

Similar to Figure S11 but with *s*=0.001. **a,** Change in inversion frequency in stochastic simulations with 10,000 individuals experiencing recessive deleterious mutations at a rate *u*=1e-08, including after inversion occurrence, all mutations having the same dominance and selection coefficients (here, *h*=0.1 and *s*=0.001). The figure displays the frequency of 10,000 inversions on an autosome and on a proto-Y chromosome, each line representing a specific inversion. **b,** Fluctuations in the mean number of mutations carried by the inverted segments in each of the 10,000 simulations, each line representing one simulation. In contrast to the Figure S11, several inversions fix on the Y chromosome before they accumulate too many mutations. Inversions take more time to fix than when $N$=1000 (Figures 3 and S10).

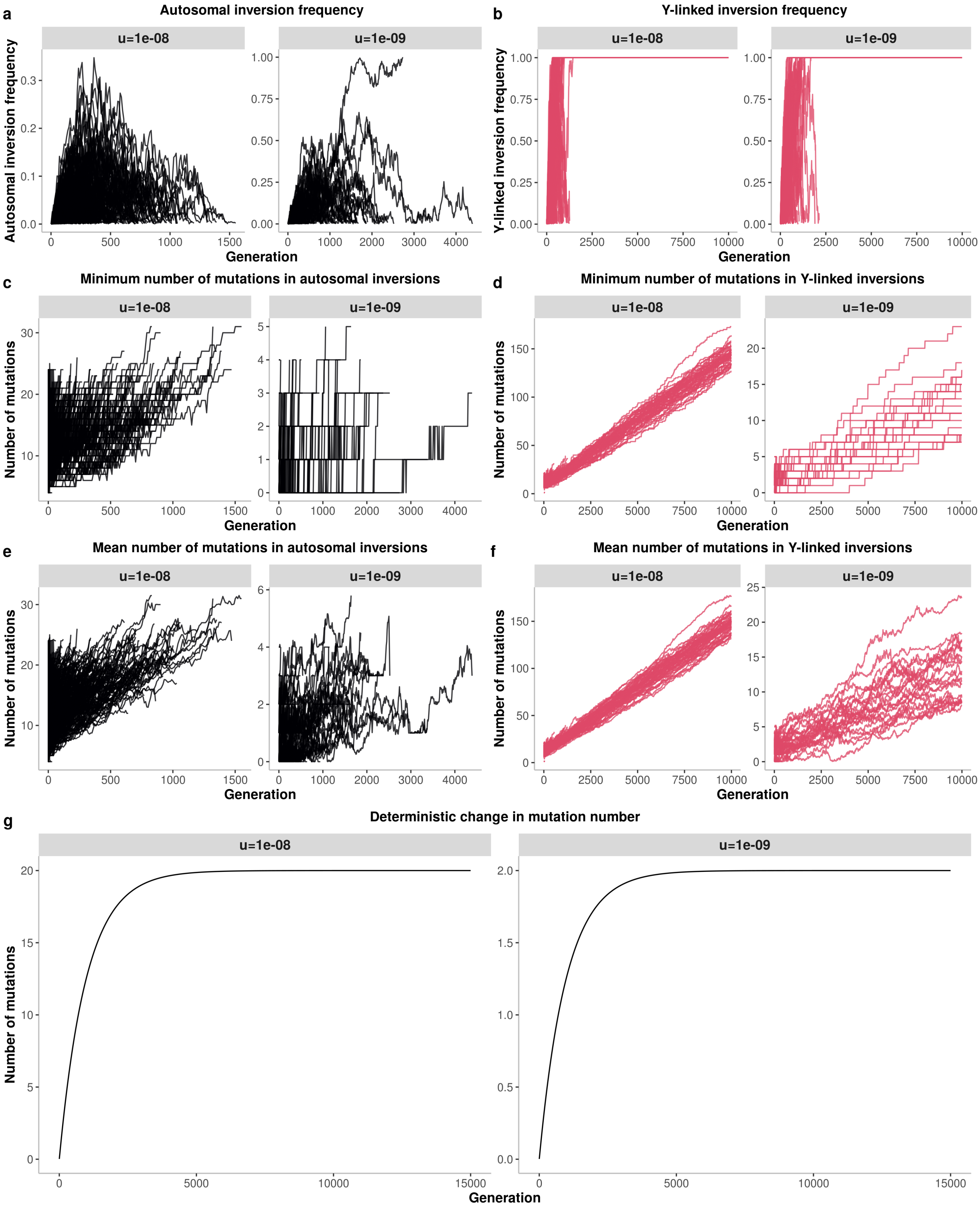


**Figure S13 | The accumulation of mutations in autosomal and sex-linked inversions in finite populations strongly depart from infinite population approximation predictions.**

**a-b,** Change in inversion frequency in stochastic simulations with *N*=1000 individuals experiencing recessive deleterious mutations at a rate *u*=1e-08 or *u*=1e-09, all mutations having the same dominance and selection coefficients (here, *h*=0.1 and *s*=0.01). The figure displays the frequency of 2 Mb inversions on an autosome and on a proto-Y chromosome. Each line represents the trajectory of one of these 10,000 simulations under each set of parameters. The lines end when no more inverted segments segregate in the population (i.e. when all inversions are lost or fixed). For *u*=1e-08, the same plots as in Figure 3a are presented but with a different scale for autosomal inversions. **c-d**, Minimum number of mutations found in inverted segments in stochastic simulations. Note the different scales of the x axes in panels c and d: as shown in panel a-b**,** no inversion segregated until the end of the simulation in the autosome (10,000 generations), thereby explaining the reduced x axis. **e-f** Same as panel c-d, but displaying the mean number of mutations in inverted segments (instead of the minimum number). For *u*=1e-08, the same plots as in figure 3b are presented (but with different scales for autosomal inversions). **g,** Deterministic change in inversion mutation number in infinite populations for a mutation rate of *u*=1e-08 or *u*=1e-09, as defined by Nei et al. (1967), and by Connallon and Olito (2021). All mutations were considered to have the same dominance and selection coefficients (here, *h*=0.1 and *s*=0.01). We considered here that inverted segments were initially mutation free. This shows that the observed accumulation of mutations in inversions on autosomes or on the Y chromosome strongly departs from the infinite population expectations.

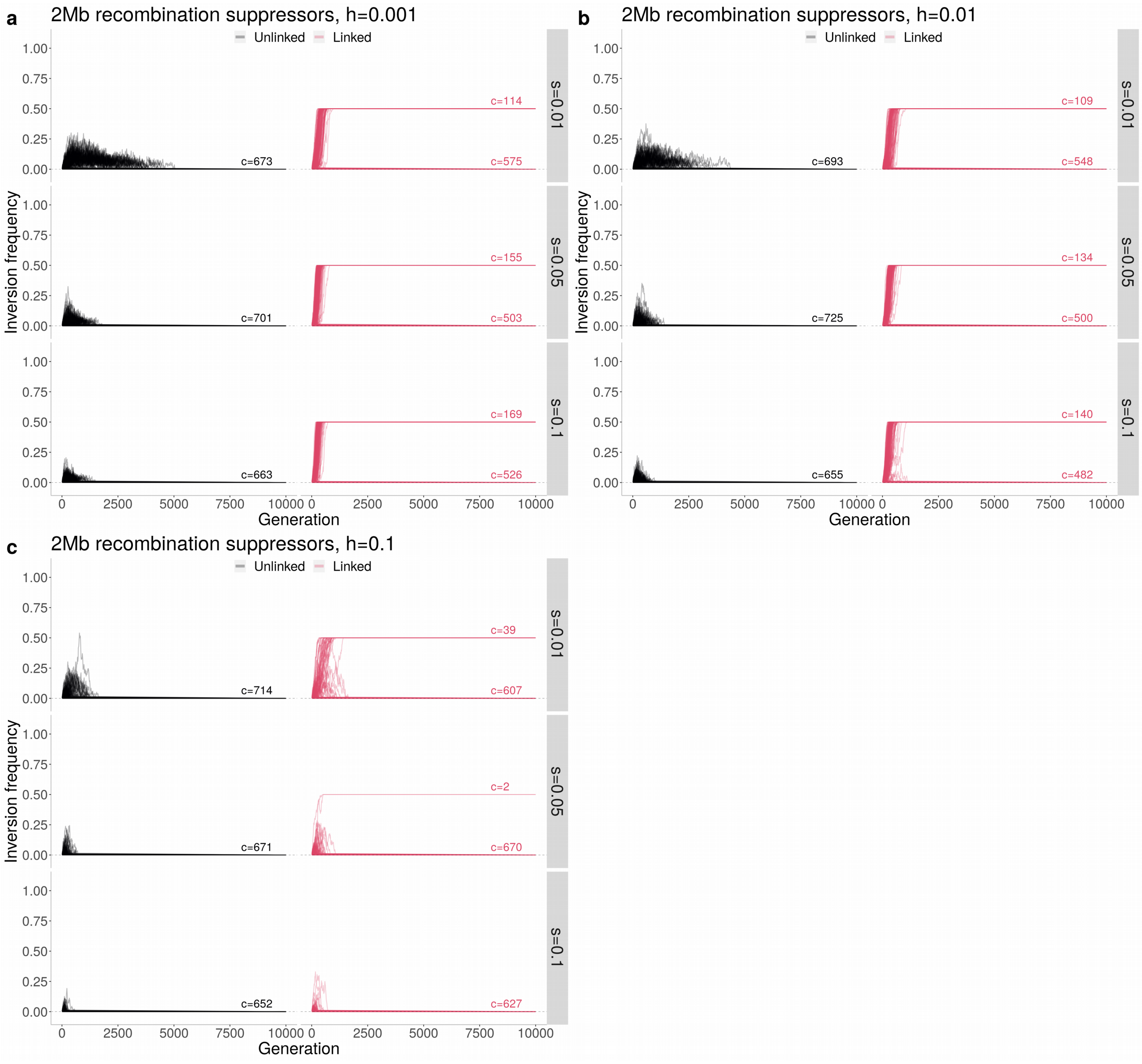


**Figure S14. Evolution of 2Mb recombination suppressors in linkage with a locus with two permanently heterozygous alleles in stochastic simulations during 10,000 generations.**

Similar to Figure 3a but with a locus with two permanently heterozygous alleles (instead of one in the case of the XY system) and with other recombination suppressors than chromosomal inversions. For each parameter combination, 10,000 recombination suppressors of 2Mb were simulated. These recombination modifiers suppress recombination within the segment in which they reside, both when heterozygous and when homozygous. These recombination suppressors were either fully linked (*r*=0.0) or fully unlinked (*r*=0.5) to a locus with two permanently heterozygous alleles; recombination suppressors fully linked to one of the two permanently heterozygous alleles can spread to a maximum frequency of 50%. Only recombination suppressors not lost after 20 generations were considered for this figure. At the end of the simulation, all recombination suppressors display either a 0.0 or a 0.5 frequency, meaning they are either lost or fixed to a given permanently heterozygous allele. The number of recombination suppressors in each state at the end of the simulation is indicated above lines (« c=... »). Populations of N=1000 individuals were simulated.

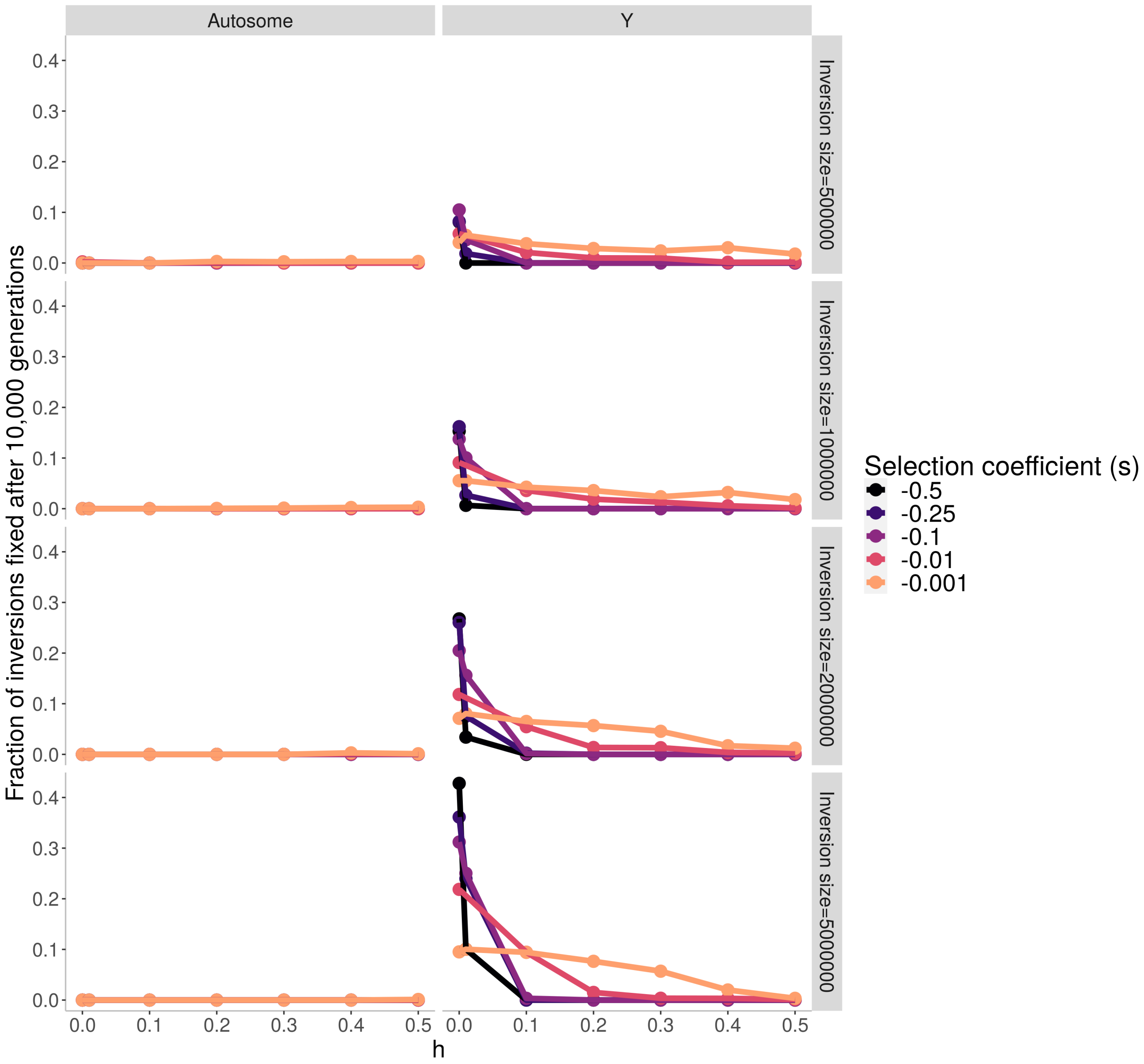


**Figure S15 | Proportions of inversions in stochastic simulations that were fixed after 10,000 generations, for different parameter value combinations and with N=1000.**
Similar to Figure 3 but considering different sizes of inversions.

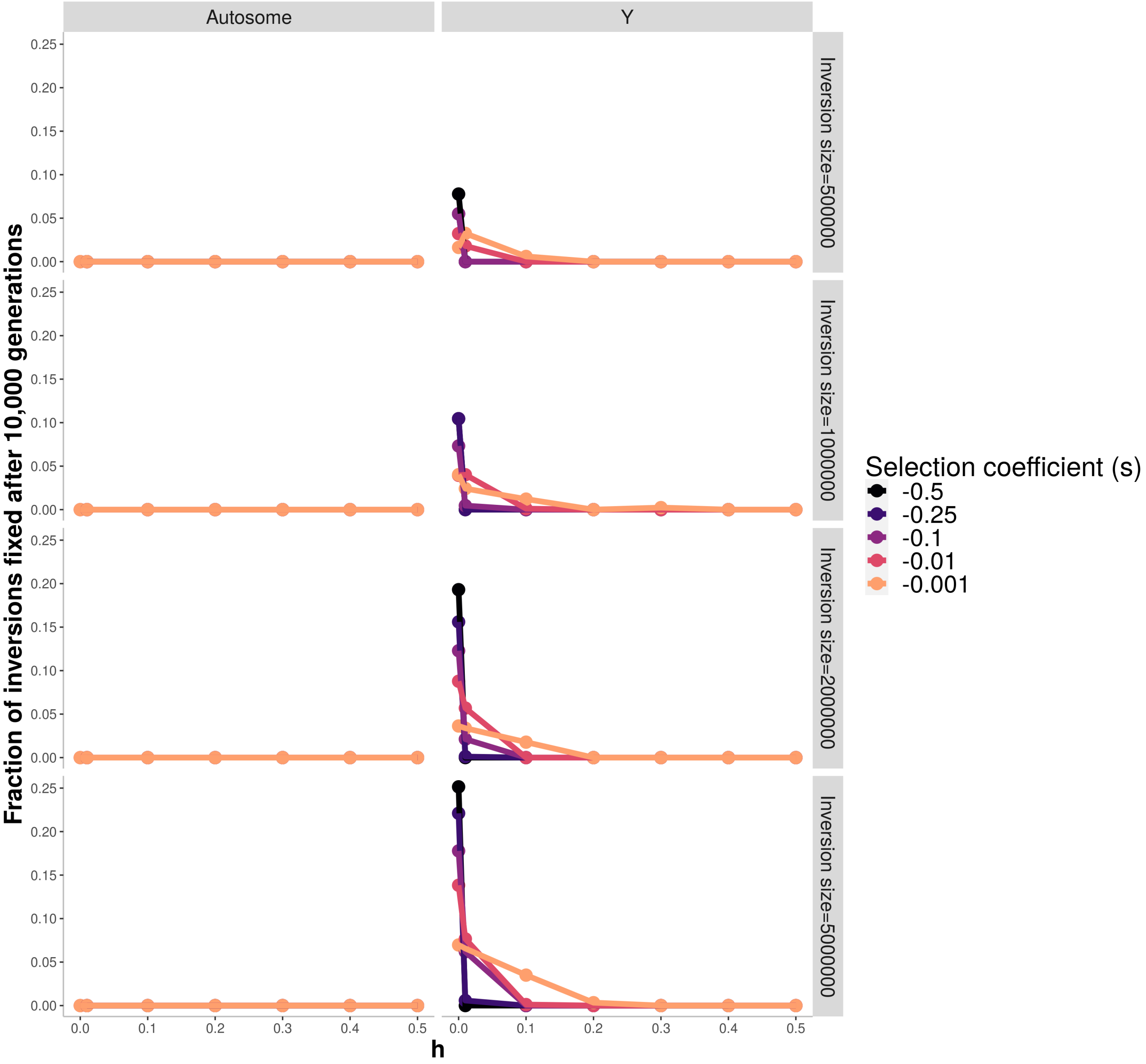


**Figure S16 | Proportions of inversions in stochastic simulations that were fixed after 10,000 generations, for different parameter value combinations with N=10,000.**

165 Similar to Figure 3 and Figure S15 but with N=10,000 and for different sizes of inversions.

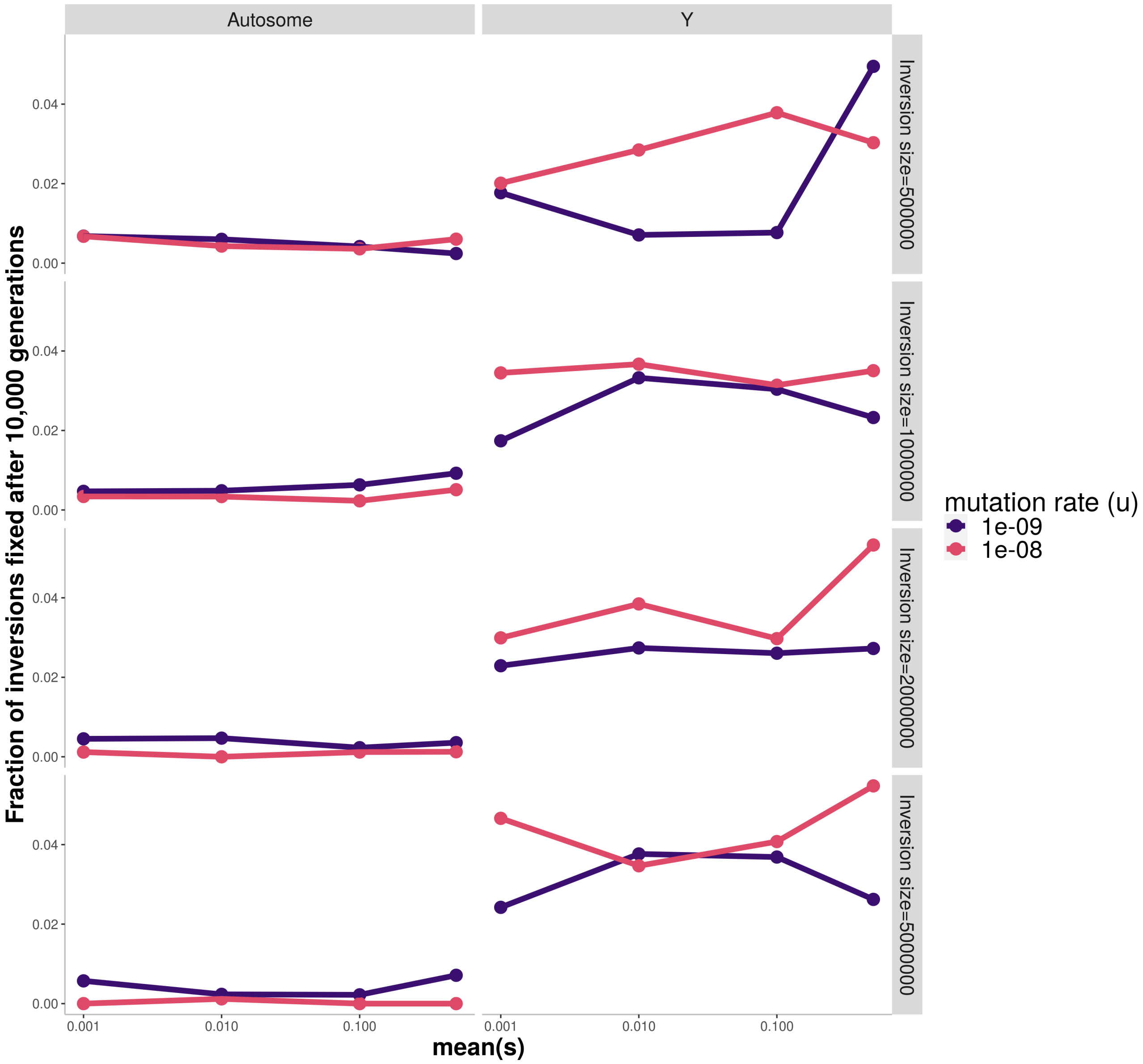


**Figure S17 | Proportions of inversions in stochastic simulations that were fixed after 10,000 generations, for different parameter value combinations with N=1000.**

Similar to Figure 3 and Figure S15 but considering that mutations segregating in the genome have their fitness effects drawn from a gamma distribution with a shape of 0.2, and their dominance coefficient *h*
170 randomly sampled among 0, 0.001, 0.01, 0.1, 0.25, 0.5 with uniform probabilities. Simulations were run considering that the mean of the selection coefficient values (mean of the gamma distribution) was either 0.001, 0.01, 0.1 or 0.5.

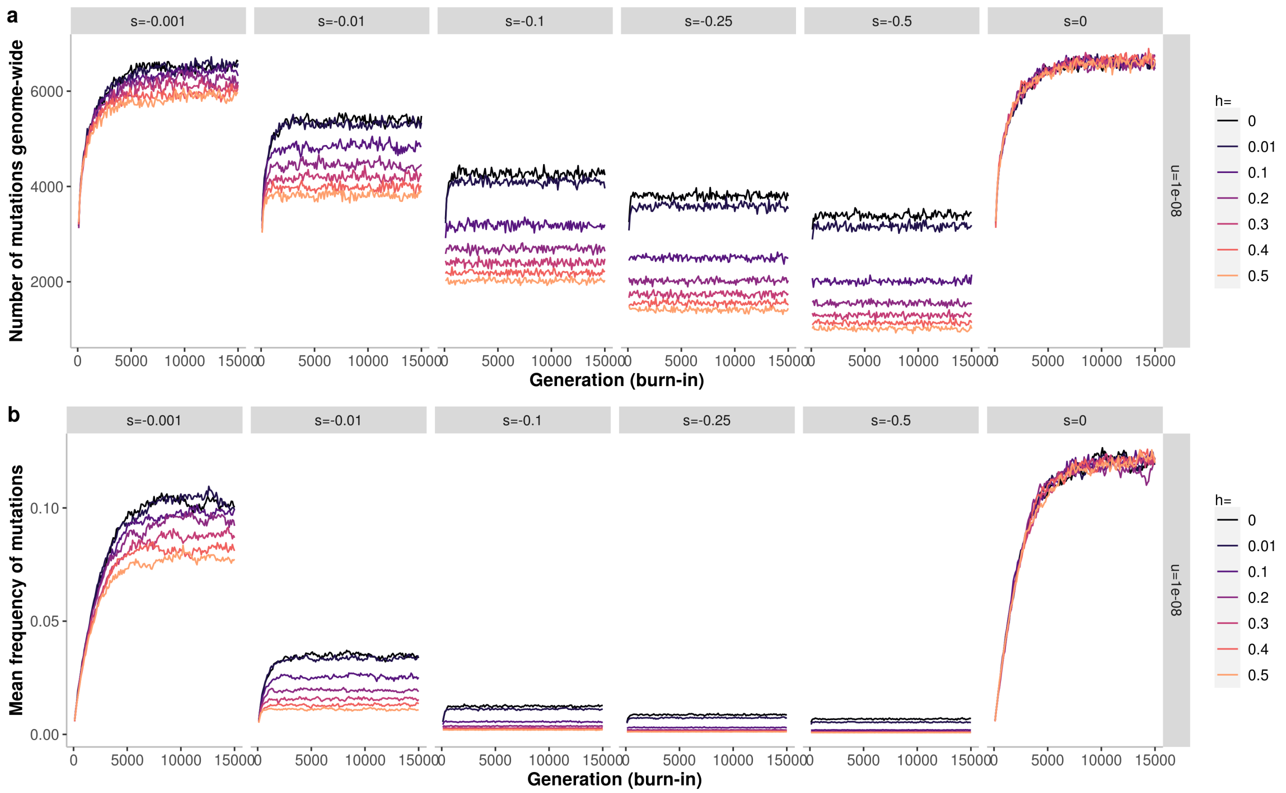


**Figure S18 | Population evolution during the burn-in phase.**

For each set of parameter values, one simulation was run and used as the initial state after a burn-in period of 15,000 generations (see methods). We display here the evolution of the population during the burn-in phase. Each trajectory represents the state of one simulation. **a,** Number of mutations segregating in the genome. **b**, Mean frequency of segregating mutations in the genome. This shows that each population had reached an equilibrium state at the end of the burn-in period.

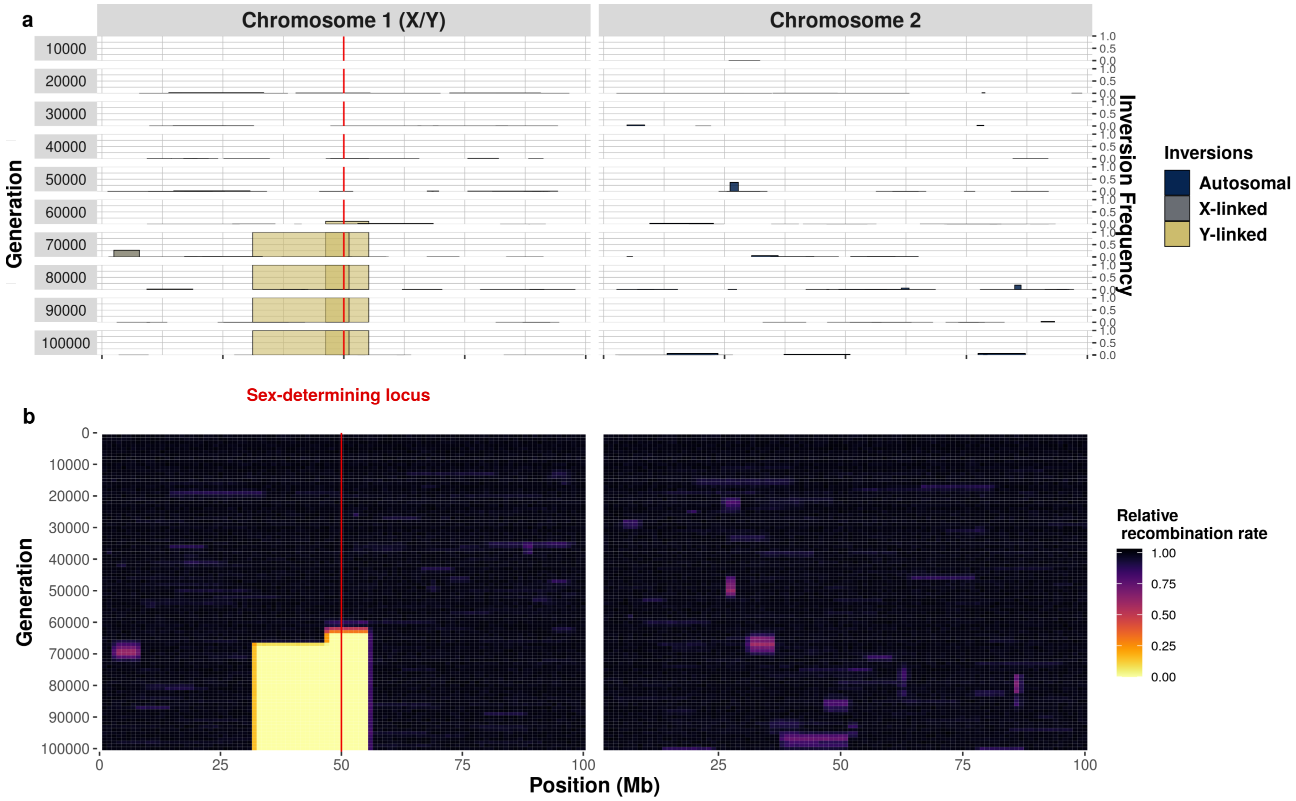


**Figure S19 | Successive accumulation of inversions around a male-determining allele in a XY system, leading to the formation of non-recombining sex chromosomes**

Similar to Figure 4 but displaying the result of another simulation and with a population of $N$=10,000 individuals. A simulation of $N$=10000 individuals, each with two pairs of 100 Mb chromosomes, during 100,000 generations. Chromosome 1 harbors an X/Y sex-determining locus at 50 Mb (individuals are XX or XY). Each generation, one inversions appears on average in the whole population, in an individual sampled uniformly at random, with the two recombination breakpoints sampled uniformly at random among $k$=100 potential breakpoints. **A,** Overview of chromosomal inversion frequency and position for 10 different generations. Square width represents inversion position and square height inversion frequency. Inversions appearing on the Y chromosome are depicted in yellow, those appearing on the X chromosomes in gray. The colors are not entirely opaque, so that regions with overlapping inversions appear darker. Loss of previously fixed inversions are due to beneficial reversion occurrence and selection. **B,** Changes in the relative rate of recombination over the entire course of the simulation. The numbers of recombination events occurring at each position (binned in 1Mb windows) are recorded at the formation of each offspring, across all homologous chromosomes in the population. To illustrate the evolution of recombination suppression between the sex chromosomes, only recombination events between the X and the Y chromosomes are shown for chromosome 1 (i.e., not the recombination events between the two X chromosomes in females). Unlike chromosome 1, chromosome 2 harbors no permanently heterozygous allele. All inversions on this chromosome suffer from homozygosity disadvantage and very few inversions therefore become fixed on chromosome 2.
