## Appendix for "Modeling the stepwise extension of recombination suppression on sex chromosomes and other supergenes through deleterious mutation sheltering"

### Appendix : Supplementary methods for "**Modeling the stepwise extension of recombination suppression on sex chromosomes and other supergenes through deleterious mutation sheltering**"

Paul Jay, Emilie Tezenas, Amandine Veber, Tatiana Giraud

#### 1 Prerequisite

We consider an infinite size, randomly mating population of diploid individuals with discrete, nonoverlapping generations. Each individual is represented by a single pair of chromosomes which carries a locus under balancing selection that may either represent a mating-type locus, a sex-determining locus or any overdominant locus, and $n$ other sites on which deleterious mutations segregate. Mutations have multiplicative effect, reducing fitness by $1 - hs$ when heterozygous, and by $1 - s$ when homozygous, $s$ and $h$ being the selection and dominance coefficients. For example, if a diploid individual has $m_1$ heterozygous sites and $m_2$ homozygous deleterious sites, with $m_1 + m_2 \leq n$, its fitness is

$$W = (1 - hs)^{m_1}(1 - s)^{m_2}.$$

Selection acts on individuals based on their genotype and individuals contribute to the next generation depending on their relative fitness. In each generation, diploid individuals are formed by randomly sampling two gametes (haploid) produced by individuals from the previous generation.

#### 2 Evolution of inversions (or any recombination suppressor acting in *cis*)

Throughout this document, we will talk about inversions but the rationale applies equally to any recombination suppressor acting in *cis* (i.e. suppressing recombination in a segment in which it resides). At a given time, an inversion appears in the population, on one chromosome. This inversion captures a segment with a number of deleterious mutations that we denote by $m$.

We are interested in studying the fate of this inversion in the population depending on $m$, $s$, $h$ and its linkage with the locus with permanently or nearly-permanently heterozygous allele(s).
Following Nei et al., (1967), we denote the presence of an inversion on a chromosome by $I$, its absence (ancestral state) by $N$.
The fitness of an individual depends on the number of mutations it carries and therefore on its genotype for the inversion.

All mutations are considered to have the same coefficient of selection and the same coefficient of dominance and the $n$ sites are supposed to be independent from one another. At mutation-selection balance, all mutations are therefore at the same frequency, denoted $q$. Hence, conditionally on an individual sequence carrying a mutation at a given site, the probability that it is paired with a sequence carrying the same mutation is q (the individual being therefore homozygous for this mutation), and the probability that it is not is $1 - q$ (the individual being therefore heterozygous).
Consider a single locus $i$ with two alleles at mutation-selection balance. The average population fitness taking only this locus into account is:

$$\begin{aligned}\overline{W_i} &= (1-q)^2 + 2q(1-q)(1-hs) + q^2(1-s) \\ &= 1 - 2q(1-q)hs - q^2 s.\end{aligned} \tag{1}$$

Since the $n$ sites are assumed independent of one another, the average fitness of an individual considering $n$ loci is:

$$\overline{W_*} = (1 - 2q(1-q)hs - q^2 s)^n, \tag{2}$$

which is the same as the average fitness of a diploid individual without inversion :

$$\overline{W_{NN}} = (1 - 2q(1-q)hs - q^2 s)^n. \tag{3}$$

Considering small $h$ and small $s$, this can be approximated by

$$\overline{W_{NN}} \approx e^{-nqsh - nqs(h+q(1-2h))}. \tag{4}$$

We can express the average fitness of an individual heterozygous for the inversion by:

$$\overline{W_{NI}} = (q(1-s) + (1-q)(1-hs))^m (q(1-hs) + 1 - q)^{n-m}, \tag{5}$$

which can again be approximated for small h and s by:

$$\overline{W_{NI}} \approx e^{-nqsh - ms(h+q(1-2h))}. \tag{6}$$

Indeed, there are $n - m$ sites where the inversion have the ancestral allele. Each of these sites have a probability $q$ of being heterozygous and $(1-q)$ of being homozygous for the ancestral allele.

Individuals homozygous for the inversion are homozygous for all mutations carried by this inversion. They have all the same fitness, which can therefore be expressed by

$$W_{II} = (1-s)^m, \tag{7}$$

and approximated by

$$W_{II} \approx e^{-sm} \tag{8}$$

when s is small.

### 3 Inversion evolutionary trajectory

We considered multiple scenarios to study the fate of an inversion depending on its linkage with an allele following various heterozygosity rule. We tracked the frequency of the inversion, considering that inversions recombine at a rate $r$ with a locus with permanently or nearly-permanently heterozygous allele(s). When $r = 0.5$, the inversion is completely unlinked with this locus and behaves as if it were on an autosome. We considered four situations:

1. The locus under balancing selection has two alleles, $X$ and $Y$, and is overdominant such that homozygotes $XX$ and $YY$ have their fitness reduced by a factor $(1 - s_2)$.

2. The locus under balancing selection has two alleles, $X$ and $Y$, and individuals are permanently heterozygotes.

3. The locus under balancing selection has three alleles, $X$, $Y$ and $Z$, and individuals are permanently heterozygotes.

4. The locus under balancing selection has two alleles, $X$ and $Y$, and individuals can be either $XX$ or $XY$ and must mate with a different genotype.

For each situation, we provide expressions for the evolution of haplotype frequencies. In each generation, genotypes frequencies are computed assuming random mating among the haplotypes. Each genotype then contributes to the next generation of haplotypes relatively to its fitness. The equations for the haplotypes frequencies are used to illustrate the deterministic change in inversion frequencies in the figures displayed in the main text and in supplementary material.

### 4 Trajectory of inversions appearing on an autosome

#### 4.1 Allelic structure

We first consider an inversion that appears on an autosome, the easiest case. The possible haplotypes are :

$$\begin{array}{rl} Haplotype & Frequency \\ N & F_N \\ I & F_I = 1 - F_N \end{array} \tag{9}$$

Then, there are three possible genotypes, which frequencies are computed assuming random mating among haplotypes.

| Genotype | Frequency | Fitness |
|---|---|---|
| $II$ | $F_I^2$ | $W_{II}$ |
| $IN$ | $2F_IF_N$ | $W_{NI}$ |
| $NN$ | $F_N^2$ | $W_{NN}$ |

The mean fitness of the population can then be computed from the genotypes frequencies as follows :

$$\overline{W} = F_N^2 W_{NN} + 2F_N F_I W_{NI} + F_I^2 W_{II}.$$

#### 4.2 Evolution of frequencies

Considering that each genotype contributes relatively to its fitness to the next generation, we compute the evolution of the frequency of the inverted haplotype after one generation :

$$F_I^{t+1} = \frac{(F_I^t)^2 W_{II} + F_I^t F_N^t W_{NI}}{\overline{W^t}}.$$

Substituing $F_N = 1 - F_I$, we obtain the variation of $F_I$ over one generation, at time $t$ :

$$\Delta F_I^t = \frac{F_I^t}{\overline{W^t}} \Big( \left(F_I^t\right)^2 (2W_{NI} - W_{II} - W_{NN}) + F_I^t (W_{II} + 2W_{NN} - 3W_{NI}) + W_{NI} - W_{NN} \Big).$$

This equation is used in Methods to derive the equilibrium frequency of an inversion that appears on an autosome.

### 5 Trajectory of inversion appearing in linkage with an overdominant locus (Fig. S8)

#### 5.1 Allelic structure

At a given locus, we consider that individuals can be either homozygotes $XX$ or $YY$, or heterozygotes $XY$, but that homozygotes suffer from a lowered fitness compared to heterozygotes (overdominance). We define $s_2$ as the selection coefficient such that homozygotes have their fitness decreased by a factor $(1 - s_2)$. For example, $W_{XI/XI} = (1 - s_2)W_{II}$. This locus being under balancing selection because of overdominance, it is hereafter referred as the "OD" locus.

Considering the evolution of an inversion I in linkage with this "OD" locus, there are four different haplotypes:

$$\begin{array}{rl} Haplotype & Frequency \\ XN & F_{XN} \\ XI & F_{XI} \\ YN & F_{YN} \\ YI & F_{YI} \end{array} \tag{10}$$

The possible genotypes and their corresponding frequencies and fitnesses are given in the following table.

| OD genotype | Inversion genotype | Frequency | Fitness |
|---|---|---|---|
| $XX$ | $II$ | $F_{XI}F_{XI}$ | $(1-s_2)W_{II}$ |
| | $IN$ | $2F_{XI}F_{XN}$ | $(1-s_2)W_{NI}$ |
| | $NN$ | $F_{XN}F_{XN}$ | $(1-s_2)W_{NN}$ |
| $XY$ | $II$ | $2F_{XI}F_{YI}$ | $W_{II}$ |
| | $IN$ | $2F_{XI}F_{YN}$ | $W_{NI}$ |
| | $NI$ | $2F_{XN}F_{YI}$ | $W_{NI}$ |
| | $NN$ | $2F_{XN}F_{YN}$ | $W_{NN}$ |
| $YY$ | $II$ | $F_{YI}F_{YI}$ | $(1-s_2)W_{II}$ |
| | $IN$ | $2F_{YI}F_{YN}$ | $(1-s_2)W_{NI}$ |
| | $NN$ | $F_{YN}F_{YN}$ | $(1-s_2)W_{NN}$ |

The frequencies are computed assuming random mating among haplotypes, and considering that $F_{XI} + F_{XN} + F_{YI} + F_{YN} = 1$.

The mean fitness of the population in a given generation can then be computed from the genotypes frequencies as follows :

$$\begin{aligned}\overline{W} &= W_{II}\Big((1-s_2)\big({F_{XI}}^2 + {F_{YI}}^2\big) + 2F_{XI}F_{YI}\Big)\\ &+ W_{NN}\Big((1-s_2)\big({F_{XN}}^2 + {F_{YN}}^2\big) + 2F_{XN}F_{YN}\Big)\\ &+ 2W_{NI}\Big((1-s_2)\big(F_{XI}F_{XN} + F_{YI}F_{YN}\big) + F_{XI}F_{YN} + F_{XN}F_{YI}\Big).\end{aligned} \tag{11}$$

#### 5.2 Evolution of frequencies

Considering a recombination rate $r$ between the overdominant locus and the inversion locus, and a generation $t$, we compute the frequency of $XI$ in the next generation :

$$\begin{aligned}F^{t+1}_{XI} &= {F^t_{XI}}^2\frac{W_{II}(1-s_2)}{\overline{W^t}} + F^t_{XI}F^t_{XN}\frac{W_{NI}(1-s_2)}{\overline{W^t}} + F^t_{XI}F^t_{YI}\frac{W_{II}}{\overline{W^t}}\\ &+ F^t_{XI}F^t_{YN}\frac{W_{NI}}{\overline{W^t}}(1-r) + F^t_{YI}F^t_{XN}\frac{W_{NI}}{\overline{W^t}}r.\end{aligned} \tag{12}$$

Re-arranging the terms, we arrive at :

$$F^{t+1}_{XI} = \frac{F^t_{XI}\Big(W_{II}\Big[(1-s_2)F^t_{XI} + F^t_{YI}\Big] + W_{NI}\Big[(1-s_2)F^t_{XN} + F^t_{YN}\Big]\Big)}{\overline{W^t}} - \frac{rW_{NI}D^t}{\overline{W^t}}, \tag{13}$$

with $D^t = F^t_{XI}F^t_{YN} - F^t_{XN}F^t_{YI}$, the linkage desequilibrium.

We can then compute the variation of $F_{XI}$ over one generation :

$$\begin{aligned}\Delta F^t_{XI} &= F^{t+1}_{XI} - F^t_{XI}\\ &= F^{t+1}_{XI} - F^t_{XI}\frac{\overline{W^t}}{\overline{W^t}}\\ &= F^t_{XI}W_{II}\frac{F^t_{YI}(1-2F^t_{XI}) + (1-s_2)\Big(F^t_{XI}(1-F^t_{XI}) - (F^t_{YI})^2\Big)}{\overline{W^t}}\\ &+ F^t_{XI}W_{NI}\frac{(1-s_2)\Big(F^t_{XN}(1-2F^t_{XI}) + 2F^t_{YI}F^t_{YN}\Big) + F^t_{YN}(1-2F^t_{XI}) - 2F^t_{XN}F^t_{YI}}{\overline{W^t}}\\ &- F^t_{XI}W^t_{NN}\frac{(1-s_2)\Big((F^t_{XN})^2 + (F^t_{YN})^2\Big) + 2F^t_{XN}F^t_{YN}}{\overline{W^t}}\\ &- \frac{rW_{NI}D^t}{\overline{W^t}}.\end{aligned} \tag{14}$$

To compute the same quantities for $F_{YI}$, we can interchange the roles of $X$ and $Y$ in the expression for $F_{XI}$, since $X$ and $Y$ have symmetrical roles:

$$F_{YI}^{t+1} = \frac{F_{YI}^t\Big(W_{II}\Big[(1-s_2)F_{YI}^t + F_{XI}^t\Big] + W_{NI}\Big[(1-s_2)F_{YN}^t + F_{XN}^t\Big]\Big)}{\overline{W^t}} + \frac{rW_{NI}D^t}{\overline{W^t}}. \quad (15)$$

From those expressions, we derive $F_{XN}^{t+1}$ by interverting $I$ and $N$ in the expression for $F_{XI}^{t+1}$, and $F_{YN}^{t+1}$ by interverting $X$ and $Y$ in the expression for $F_{XN}^{t+1}$ :

$$\begin{aligned} F_{XN}^{t+1} &= \frac{F_{XN}^t\Big(W_{NN}\Big[(1-s_2)F_{XN}^t + F_{YN}^t\Big] + W_{NI}\Big[(1-s_2)F_{XI}^t + F_{YI}^t\Big]\Big)}{\overline{W^t}} + \frac{rW_{NI}D^t}{\overline{W^t}}, \\ F_{YN}^{t+1} &= \frac{F_{YN}^t\Big(W_{NN}\Big[(1-s_2)F_{YN}^t + F_{XN}^t\Big] + W_{NI}\Big[(1-s_2)F_{YI}^t + F_{XI}^t\Big]\Big)}{\overline{W^t}} - \frac{rW_{NI}D^t}{\overline{W^t}}. \end{aligned} \quad (16)$$

Figure S8 shows a numerical solving of these equations.

### 6 Trajectory of inversions in linkage with a permanently heterozygous locus with two alleles (Figs. S5-6)

#### 6.1 Allelic structure

At a given locus with two alleles, X and Y, we suppose that individuals are all heterozygotes $XY$.

Considering an inversion I segregating in this population, there are still the same four possible haplotypes in the population:

$$\begin{array}{rl} Haplotype & Frequency \\ XN & F_{XN} \\ XI & F_{XI} \\ YN & F_{YN} \\ YI & F_{YI} \end{array} \quad (17)$$

Since a diploid genotype can only be formed by two chromosomes that have different alleles (X or Y), the frequencies of $X$-chromosomes and of $Y$-chromosomes are equal. We therefore consider independently the frequency of inversions in proto-X chromosomes and in proto-Y chromosomes, and we can make the assumption that $F_{XN} + F_{XI} = 1$ and $F_{YN} + F_{YI} = 1$ to ease the calculations.

There are four possible genotypes, which frequencies are computed assuming random mating among haplotypes.

| Genotype | Frequency | Fitness |
|---|---|---|
| $XN/YN$ | $F_{XN}F_{YN}$ | $W_{NN}$ |
| $XI/YN$ | $F_{XI}F_{YN}$ | $W_{NI}$ |
| $XN/YI$ | $F_{XN}F_{YI}$ | $W_{NI}$ |
| $XI/YI$ | $F_{XI}F_{YI}$ | $W_{II}$ |

The mean fitness of the population is thus

$$\overline{W} = F_{XN}F_{YN}W_{NN} + \big(F_{XI}F_{YN} + F_{XN}F_{YI}\big)W_{NI} + F_{XI}F_{YI}W_{II}.$$

#### 6.2 Evolution of frequencies

Considering a recombination rate $r$ between the permanently heterozygous locus and the inversion locus, and a generation $t$, we can compute the frequency of the haplotype XI in the next generation:

$$\begin{aligned} F_{XI}^{t+1} &= \frac{F_{XI}^t F_{YI}^t W_{II} + F_{XI}^t F_{YN}^t W_{NI}(1-r) + F_{XN}^t F_{YI}^t W_{NI} r}{\overline{W^t}} \\ &= \frac{F_{XI}^t (F_{YI}^t W_{II} + F_{YN}^t W_{NI}) - r W_{NI}(F_{XI}^t F_{YN}^t - F_{XN}^t F_{YI}^t)}{\overline{W^t}} \\ &= \frac{F_{XI}^t W_{XI}^* - r W_{NI} D^t}{\overline{W^t}}, \end{aligned} \tag{18}$$

with $D^t = F_{XI}^t F_{YN}^t - F_{XN}^t F_{YI}^t$, the linkage disequilibrium, and $W_{XI}^* = F_{YN} W_{NI} + F_{YI} W_{II}$, the marginal fitness of haplotype $XI$.

The change in frequency in each generation is then

$$\begin{aligned} \Delta F_{XI}^t &= F_{XI}^{t+1} - F_{XI}^t \\ &= \frac{F_{XI}^t W_{XI}^* - r W_{NI} D^t}{\overline{W^t}} - \frac{F_{XI}^t \overline{W^t}}{\overline{W^t}} \\ &= \frac{F_{XI}^t (W_{XI}^* - \overline{W^t}) - r W_{NI} D^t}{\overline{W^t}} \\ &= \frac{F_{XI}^t F_{XN}^t (W_{XI}^* - W_{XN}^*) - r W_{NI} D^t}{\overline{W^t}} \\ &= \frac{F_{XI}^t (1 - F_{XI}^t)(W_{XI}^* - W_{XN}^*) - r W_{NI} D^t}{\overline{W^t}}. \end{aligned} \tag{19}$$

As before, we can determine the frequencies of $XN$, $YI$ or $YN$ in generation $t+1$ by interchanging the roles of X and Y, or of I and N in the equation for $F_{XI}^{t+1}$ to obtain

$$\begin{aligned} F_{XN}^{t+1} &= \frac{F_{XN}^t (F_{YN}^t W_{NN} + F_{YI}^t W_{NI}) + r W_{NI} D^t}{\overline{W^t}}, \\ F_{YN}^{t+1} &= \frac{F_{YN}^t (F_{XN}^t W_{NN} + F_{XI}^t W_{NI}) - r W_{NI} D^t}{\overline{W^t}}, \\ F_{YI}^{t+1} &= \frac{F_{YI}^t (F_{XI}^t W_{II} + F_{XI}^t W_{NI}) + r W_{NI} D^t}{\overline{W^t}}. \end{aligned} \tag{20}$$

We observe that these equations are the same as in the overdominant case when $s_2 = 1$, *i.e.* when no homozygous are formed.

### 7 Trajectory of inversions in linkage with a permanently heterozygous locus with three alleles (Fig. S7)

#### 7.1 Allelic structure

At a given locus with three alleles, $X$, $Y$ and $Z$, we consider that individuals are all heterozygotes $XY$, $XZ$ or $YZ$.

There are six possible haplotypes :

$$\begin{array}{rl} Haplotype & Frequency \\ XN & F_{XN} \\ XI & F_{XI} \\ YN & F_{YN} \\ YI & F_{YI} \\ ZN & F_{ZN} \\ ZI & F_{ZI} \end{array} \tag{21}$$

The derivation of the frequency of each genotype is a bit more challenging than in the case of two permanently heterozygous alleles.

We denote the frequency of chromosomes with allele $X$ among all chromosomes by $F_X$. We thus have $F_X = F_{XI} + F_{XN}$, and likewise for $F_Y$ and $F_Z$. We also have $F_X + F_Y + F_Z = 1$.

We can compute the frequency of each genotype (e.g $F_{XIYN}$, the frequency of the genotype formed by a haplotype $XI$ and a haplotype $YN$) assuming random mating. For instance, to compute $F_{XIYN}$, we have first to pick one of the two haplotypes among all haplotypes (either $XI$ with probability $F_{XI}$ or $YN$ with probability $F_{YN}$), and then pick the second among the remaining compatible haplotypes. If $XI$ was picked first, the second haplotype has to carry either the $Y$ allele or the $Z$ allele to ensure the heterozygosity. Among those compatible haplotypes, the $YN$ haplotype has a probability of $\frac{F_{YN}}{F_Y+F_Z}$ to be picked. We have thus :

$$\begin{aligned}
F_{XIYN} &= \mathbb{P}(\text{choose } XI \text{ first})\mathbb{P}(\text{choose } YN \text{ after a } XI) + \mathbb{P}(\text{choose } YN \text{ first})\mathbb{P}(\text{choose } XI \text{ after a } YN) \\
&= \mathbb{P}(\text{choose } XI \text{ first})\mathbb{P}(\text{choose } YN \text{ among the } Y \text{ and } Z \text{ chrom.}) \\
&\qquad + \mathbb{P}(\text{choose } YN \text{ first})\mathbb{P}(\text{choose } XI \text{ among the } X \text{ and } Z \text{ chrom.}) \\
&= F_{XI}\frac{F_{YN}}{F_Y + F_Z} + F_{YN}\frac{F_{XI}}{F_X + F_Z} \\
&= F_{XI}\frac{F_{YN}}{1 - F_X} + F_{YN}\frac{F_{XI}}{1 - F_Y}.
\end{aligned}$$

We reduce the fractions to a common denominator, and use the equation $F_X + F_Y + F_Z = 1$ to obtain :

$$\begin{aligned}
F_{XIYN} &= F_{XI}F_{YN}\left(\frac{1 - F_Y}{(1 - F_X)(1 - F_Y)} + \frac{1 - F_X}{(1 - F_X)(1 - F_Y)}\right) \\
&= F_{XI}F_{YN}\left(\frac{1 + 1 - F_Y - F_X}{(1 - F_X)(1 - F_Y)}\right) \\
&= F_{XI}F_{YN}\frac{1 + F_Z}{(1 - F_X)(1 - F_Y)}.
\end{aligned}$$

The possible genotypes and their corresponding frequencies are therefore:

| OD genotype | Inversion genotype | Frequency | Fitness |
|---|---|---|---|
| $XY$ | $II$ | $F_{XI}F_{YI}\frac{1+F_Z}{(1-F_X)(1-F_Y)}$ | $W_{II}$ |
| | $IN$ | $F_{XI}F_{YN}\frac{1+F_Z}{(1-F_X)(1-F_Y)}$ | $W_{IN}$ |
| | $NI$ | $F_{XN}F_{YI}\frac{1+F_Z}{(1-F_X)(1-F_Y)}$ | $W_{IN}$ |
| | $NN$ | $F_{XN}F_{YN}\frac{1+F_Z}{(1-F_X)(1-F_Y)}$ | $W_{NN}$ |
| $XZ$ | $II$ | $F_{XI}F_{ZI}\frac{1+F_Y}{(1-F_X)(1-F_Z)}$ | $W_{II}$ |
| | $IN$ | $F_{XI}F_{ZN}\frac{1+F_Y}{(1-F_X)(1-F_Z)}$ | $W_{IN}$ |
| | $NI$ | $F_{XN}F_{ZI}\frac{1+F_Y}{(1-F_X)(1-F_Z)}$ | $W_{IN}$ |
| | $NN$ | $F_{XN}F_{ZN}\frac{1+F_Y}{(1-F_X)(1-F_Z)}$ | $W_{NN}$ |
| $YZ$ | $II$ | $F_{YI}F_{ZI}\frac{1+F_X}{(1-F_Y)(1-F_Z)}$ | $W_{II}$ |
| | $IN$ | $F_{YI}F_{ZN}\frac{1+F_X}{(1-F_Y)(1-F_Z)}$ | $W_{IN}$ |
| | $NI$ | $F_{YN}F_{ZI}\frac{1+F_X}{(1-F_Y)(1-F_Z)}$ | $W_{IN}$ |
| | $NN$ | $F_{YN}F_{ZN}\frac{1+F_X}{(1-F_Y)(1-F_Z)}$ | $W_{NN}$ |

The mean fitness of the population is thus

$$\begin{aligned}\overline{W} =& F_{XIYI}W_{II} + F_{XIYN}W_{NI} + F_{XNYI}W_{NI} + F_{XNYN}W_{NN} + \\ & F_{XIZI}W_{II} + F_{XIZN}W_{NI} + F_{XNZI}W_{NI} + F_{XNZN}W_{NN} + \\ & F_{YIZI}W_{II} + F_{YIZN}W_{NI} + F_{YNZI}W_{NI} + F_{YNZN}W_{NN}.\end{aligned} \tag{22}$$

#### 7.2 Evolution of frequencies

Considering a recombination rate $r$ between the permanently heterozygous locus and the inversion, and a generation $t$, we can compute the frequency of each haplotype in the next generation. We write $F_X^t := F_{XN}^t + F_{XI}^t$ (likewise for $F_Y^t$ and $F_Z^t$) and set $\widehat{F}^t := (1-F_X^t)(1-F_Y^t)(1-F_Z^t)$. We have

$$\begin{aligned}F_{XI}^{t+1} =& \frac{W_{II}}{2\overline{W}^t}\frac{F_{XI}^t}{\widehat{F}^t}\Big[F_{YI}^t\left(1-(F_Z^t)^2\right) + F_{ZI}^t\left(1-(F_Y^t)^2\right)\Big] \\ &+ \frac{W_{IN}}{2\overline{W}^t}\frac{F_{XI}^t}{\widehat{F}^t}\Big[F_{YN}^t\left(1-(F_Z^t)^2\right) + F_{ZN}^t\left(1-(F_Y^t)^2\right)\Big] \\ &+ r\frac{W_{IN}}{2\overline{W}^t}\left(\frac{F_{XN}^t}{\widehat{F}^t}\Big[F_{YI}^t\left(1-(F_Z^t)^2\right) + F_{ZI}^t\left(1-(F_Y^t)^2\right)\Big] - \frac{F_{XI}^t}{\widehat{F}^t}\Big[F_{YN}^t\left(1-(F_Z^t)^2\right) + F_{ZN}^t\left(1-(F_Y^t)^2\right)\Big]\right)\end{aligned} \tag{23}$$

$$\begin{aligned}F_{XN}^{t+1} =& \frac{W_{NN}}{2\overline{W}^t}\frac{F_{XN}^t}{\widehat{F}^t}\Big[F_{YN}^t\left(1-(F_Z^t)^2\right) + F_{ZN}^t\left(1-(F_Y^t)^2\right)\Big] \\ &+ \frac{W_{IN}}{2\overline{W}^t}\frac{F_{XN}^t}{\widehat{F}^t}\Big[F_{YI}^t\left(1-(F_Z^t)^2\right) + F_{ZI}^t\left(1-(F_Y^t)^2\right)\Big] \\ &+ r\frac{W_{IN}}{2\overline{W}^t}\left(\frac{F_{XI}^t}{\widehat{F}^t}\Big[F_{YN}^t\left(1-(F_Z^t)^2\right) + F_{ZN}^t\left(1-(F_Y^t)^2\right)\Big] - \frac{F_{XN}^t}{\widehat{F}^t}\Big[F_{YI}^t\left(1-(F_Z^t)^2\right) + F_{ZI}^t\left(1-(F_Y^t)^2\right)\Big]\right)\end{aligned} \tag{24}$$

$$\begin{aligned}F_{YI}^{t+1} =& \frac{W_{II}}{2\overline{W}^t}\frac{F_{YI}^t}{\widehat{F}^t}\Big[F_{XI}^t\left(1-(F_Z^t)^2\right) + F_{ZI}^t\left(1-(F_X^t)^2\right)\Big] \\ &+ \frac{W_{IN}}{2\overline{W}^t}\frac{F_{YI}^t}{\widehat{F}^t}\Big[F_{XN}^t\left(1-(F_Z^t)^2\right) + F_{ZN}^t\left(1-(F_X^t)^2\right)\Big] \\ &+ r\frac{W_{IN}}{2\overline{W}^t}\left(\frac{F_{YN}^t}{\widehat{F}^t}\Big[F_{XI}^t\left(1-(F_Z^t)^2\right) + F_{ZI}^t\left(1-(F_X^t)^2\right)\Big] - \frac{F_{YI}^t}{\widehat{F}^t}\Big[F_{XN}^t\left(1-(F_Z^t)^2\right) + F_{ZN}^t\left(1-(F_X^t)^2\right)\Big]\right)\end{aligned} \tag{25}$$

$$\begin{aligned}F_{YN}^{t+1} =& \frac{W_{NN}}{2\overline{W}^t}\frac{F_{YN}^t}{\widehat{F}^t}\Big[F_{XN}^t\left(1-(F_Z^t)^2\right) + F_{ZN}^t\left(1-(F_X^t)^2\right)\Big] \\ &+ \frac{W_{IN}}{2\overline{W}^t}\frac{F_{YN}^t}{\widehat{F}^t}\Big[F_{XI}^t\left(1-(F_Z^t)^2\right) + F_{ZI}^t\left(1-(F_X^t)^2\right)\Big] \\ &+ r\frac{W_{IN}}{2\overline{W}^t}\left(\frac{F_{YI}^t}{\widehat{F}^t}\Big[F_{XN}^t\left(1-(F_Z^t)^2\right) + F_{ZN}^t\left(1-(F_X^t)^2\right)\Big] - \frac{F_{YN}^t}{\widehat{F}^t}\Big[F_{XI}^t\left(1-(F_Z^t)^2\right) + F_{ZI}^t\left(1-(F_X^t)^2\right)\Big]\right)\end{aligned} \tag{26}$$

$$\begin{aligned}F_{ZI}^{t+1} =& \frac{W_{II}}{2\overline{W}^t}\frac{F_{ZI}^t}{\widehat{F}^t}\Big[F_{XI}^t\left(1-(F_Y^t)^2\right) + F_{YI}^t\left(1-(F_X^t)^2\right)\Big] \\ &+ \frac{W_{IN}}{2\overline{W}^t}\frac{F_{ZI}^t}{\widehat{F}^t}\Big[F_{XN}^t\left(1-(F_Y^t)^2\right) + F_{YN}^t\left(1-(F_X^t)^2\right)\Big] \\ &+ r\frac{W_{IN}}{2\overline{W}^t}\left(\frac{F_{ZN}^t}{\widehat{F}^t}\Big[F_{XI}^t\left(1-(F_Y^t)^2\right) + F_{YI}^t\left(1-(F_X^t)^2\right)\Big] - \frac{F_{ZI}^t}{\widehat{F}^t}\Big[F_{XN}^t\left(1-(F_Y^t)^2\right) + F_{YN}^t\left(1-(F_X^t)^2\right)\Big]\right)\end{aligned} \tag{27}$$

$$
\begin{aligned}
F_{ZN}^{t+1} = {} & \frac{W_{NN}}{2\overline{W^t}} \frac{F_{ZN}^t}{\widehat{F}^t} \left[ F_{XN}^t \left(1 - (F_Y^t)^2\right) + F_{YN}^t \left(1 - (F_X^t)^2\right) \right] \\
& + \frac{W_{IN}}{2\overline{W^t}} \frac{F_{ZN}^t}{\widehat{F}^t} \left[ F_{XI}^t \left(1 - (F_Y^t)^2\right) + F_{YI}^t \left(1 - (F_X^t)^2\right) \right] \\
& + r \frac{W_{IN}}{2\overline{W^t}} \left( \frac{F_{ZI}^t}{\widehat{F}^t} \left[ F_{XN}^t \left(1 - (F_Y^t)^2\right) + F_{YN}^t \left(1 - (F_X^t)^2\right) \right] - \frac{F_{ZN}^t}{\widehat{F}^t} \left[ F_{XI}^t \left(1 - (F_Y^t)^2\right) + F_{YI}^t \left(1 - (F_X^t)^2\right) \right] \right)
\end{aligned} \tag{28}
$$

The expression for the change in frequency over one generation can be obtained from the above formulae. Since the expression is complex and not particularly informative, we do not detail it here.

If we set $F_{ZI} = F_{ZN} = 0$, we obtain the same equations as in the case of two alleles for the permanent heterozygous locus.

### 8 Trajectory of inversions in linkage with an XY mammal-like sex-determining locus (Figs. 2, 3 & S4)

#### 8.1 Allelic structure

At a given locus with two alleles, X and Y, we suppose that individuals can be either $XX$ or $XY$ and must mate with a different genotype.

There are still the same four possible haplotypes in the population, but, following Clarke (1987), it is useful to distinguish the frequency of inverted and non-inverted segments in proto-X chromosomes in gametes produced by males ($F_{XIm}$) and by females ($F_{XIf}$).

| Haplotype | Frequency in male | Frequency in female |
|---|---|---|
| XN | $F_{XNm}$ | $F_{XNf}$ |
| XI | $F_{XIm}$ | $F_{XIf}$ |
| YN | $F_{YN}$ | 0 |
| YI | $F_{YI}$ | 0 |

We consider independently the frequency of inversions in proto-X chromosomes and proto-Y chromosomes so that we can assume $F_{XNf} + F_{XIf} = 1$, $F_{XNm} + F_{XIm} = 1$ and $F_{YN} + F_{YI} = 1$ to ease the calculations.

There are seven possible genotypes which frequencies can be computed assuming random mating.

| Genotype | Frequency | Fitness |
|---|---|---|
| $XN/XN$ | $F_{XNm}F_{XNf}$ | $W_{NN}$ |
| $XN/XI$ | $F_{XNm}F_{XIf} + F_{XNf}F_{XIm}$ | $W_{NI}$ |
| $XI/XI$ | $F_{XIm}F_{XIf}$ | $W_{II}$ |
| $XN/YN$ | $F_{XNf}F_{YN}$ | $W_{NN}$ |
| $XI/YN$ | $F_{XIf}F_{YN}$ | $W_{NI}$ |
| $XN/YI$ | $F_{XNf}F_{YI}$ | $W_{NI}$ |
| $XI/YI$ | $F_{XIf}F_{YI}$ | $W_{II}$ |

We consider independently the fitness of males ($\overline{W_m}$) and females ($\overline{W_f}$). These quantities are given by

$$\overline{W_m} = F_{XNf}F_{YN}W_{NN} + F_{XNf}F_{YI}W_{NI} + F_{XIf}F_{YN}W_{NI} + F_{XIf}F_{YI}W_{II}$$

and

$$\overline{W_f} = F_{XNf}F_{XNm}W_{NN} + F_{XNm}F_{XIf}W_{NI} + F_{XNf}F_{XIm}W_{NI} + F_{XIf}F_{XIm}W_{II}.$$

##### 8.2 Evolution of frequencies

Considering a recombination rate $r$ between the sex-determining locus and the inversion, and a generation $t$, we can follow the frequency of inverted and non-inverted segments in proto-X chromosomes independently in gametes produced by males ($F_{XIm}$) and by females ($F_{XIf}$):

$$
\begin{aligned}
F_{XIm}^{t+1} &= \frac{F_{XIf}^t F_{YI}^t W_{II} + F_{XIf}^t F_{YN}^t W_{NI}(1-r) + F_{XNf}^t F_{YI}^t W_{NI} r}{\overline{W_m^t}} \\
&= \frac{F_{XIf}^t \left(W_{II} F_{YI}^t + W_{NI} F_{YN}^t\right) + r W_{NI} D^t}{\overline{W_m^t}}, \\
F_{XNm}^{t+1} &= \frac{F_{XNf}^t F_{YN}^t W_{NN} + F_{XNf}^t F_{YI}^t W_{NI}(1-r) + F_{XIf}^t F_{YN}^t W_{NI} r}{\overline{W_m^t}} \\
&= \frac{F_{XNf}^t \left(W_{NN} F_{YN}^t + W_{NI} F_{YI}^t\right) - r W_{NI} D^t}{\overline{W_m^t}}, \\
F_{XIf}^{t+1} &= \frac{F_{XIm}^t F_{XIf}^t W_{II} + \frac{1}{2} W_{NI} [F_{XIm}^t F_{XNf}^t + F_{XIf}^t F_{XNm}^t]}{\overline{W_f^t}}, \\
F_{XNf}^{t+1} &= \frac{F_{XNm}^t F_{XNf}^t W_{NN} + \frac{1}{2} W_{NI} [F_{XNm}^t F_{XIf}^t + F_{XNf}^t F_{XIm}^t]}{\overline{W_f^t}},
\end{aligned}
\tag{29}
$$

with $D^t = F_{XNf}^t F_{YI}^t - F_{XIf}^t F_{YN}^t$.

Since two third of the X chromosomes are in females and one third are in male, the frequency of the inversion in proto-X chromosomes in the whole population can be obtained by:

$$
F_{XI} = \frac{2F_{XIf}}{3} + \frac{F_{XIm}}{3}.
$$

The variation in the frequency of $XI$ over one generation is then :

$$
\begin{aligned}
\Delta F_{XI}^t &= F_{XI}^{t+1} - F_{XI}^t \\
&= \frac{2}{3}\left(F_{XIf}^{t+1} - F_{XIf}^t\right) + \frac{1}{3}\left(F_{XIm}^{t+1} - F_{XIm}^t\right) \\
&= \frac{2}{3}\frac{1}{\overline{W_f^t}}\Big[F_{XIf}^t F_{XNf}^t \left(W_{II} F_{XIm}^t - W_{NN} F_{XNm}^t\right) \\
&\qquad + W_{NI}\left(\frac{1}{2} - F_{XIf}^t\right)\left(F_{XIm}^t F_{XNf}^t + F_{XIf}^t F_{XNm}^t\right)\Big] \\
&+ \frac{1}{3}\frac{1}{\overline{W_m^t}}\Big[W_{II} F_{YI}^t F_{XIf}^t F_{XNm}^t - W_{NN} F_{XIm}^t F_{XNf}^t F_{YN}^t \\
&\qquad + W_{NI}\left(F_{YN}^t F_{XIf}^t F_{XNm}^t - F_{XIm}^t F_{XNf}^t F_{YI}^t\right) + r W_{NI} D^t\Big]
\end{aligned}
\tag{30}
$$

The changes in frequency of inverted and non-inverted segments in proto-Y chromosomes are calculated as before:

$$
\begin{aligned}
F_{YI}^{t+1} &= \frac{F_{YI}^t (F_{XIf}^t W_{II} + F_{XNf}^t W_{NI}) - r W_{NI} D^t}{\overline{W_m^t}}, \\
F_{YN}^{t+1} &= \frac{F_{YN}^t (F_{XNf}^t W_{NN} + F_{XIf}^t W_{NI}) + r W_{NI} D^t}{\overline{W_m^t}}.
\end{aligned}
\tag{31}
$$

#### 9 Deterministic assessment of inversion fate

The fate of inversions can be determined as a function of their fitness. As summarized in the Methods sections, inversions linked to a Y allele or on autosomes can initially increase in frequency only if heterozygotes for the inversion have a better fitness than homozygotes for the absence of inversion (i.e. $W_{NI} > W_{NN}$). This condition is sufficient for a Y-linked inversion to go to fixation, whereas on autosomes, inversion can go to fixation only if homozygotes for the inversion must have a better fitness than heterozygotes for the inversion ($W_{II} > W_{NI}$). We can easily derive the conditions on the number of mutations carried by the inversion ($m$) for these situations to occur. We have

$$\begin{aligned}
& W_{II} > W_{NI} \\
\text{if and only if} \quad & (1-s)^m > (q(1-s)+(1-q)(1-hs))^m (q(1-hs)+1-q)^{n-m}, \\
\text{i.e.} \quad & \left[\frac{(1-s)\big(q(1-hs)+1-q\big)}{q(1-s)+(1-q)(1-hs)}\right]^m > [q(1-hs)+1-q]^n, \\
\text{i.e.} \quad & m \ln\left[\frac{(1-s)\big(q(1-hs)+1-q\big)}{q(1-s)+(1-q)(1-hs)}\right] > n \ln[q(1-hs)+1-q].
\end{aligned}$$

Since $1-s < 1-hs < 1$, we have $(1-s)\big(q(1-hs)+1-q\big) = q(1-s)(1-hs)+(1-q)(1-q) < q(1-s)+(1-q)(1-hq)$, and thus $\ln\left[\frac{(1-s)\big(q(1-hs)+1-q\big)}{q(1-s)+(1-q)(1-hs)}\right] < 0$.

Therefore,

$$W_{II} > W_{NI} \quad \text{if and only if} \quad m < n \times \frac{\ln[q(1-hs)+1-q]}{\ln\left[\frac{(1-s)\big(q(1-hs)+1-q\big)}{q(1-s)+(1-q)(1-hs)}\right]} = n \times \beta_1(q,h,s).$$

Similarly, we have

$$\begin{aligned}
& W_{NI} > W_{NN} \\
\text{if and only if} \quad & (q(1-s)+(1-q)(1-hs))^m (q(1-hs)+1-q)^{n-m} > (1-2q(1-q)hs-q^2 s)^n, \\
\text{i.e.} \quad & \left[\frac{q(1-s)+(1-q)(1-hs)}{q(1-hs)+1-q}\right]^m > \left[\frac{1-2q(1-q)hs-q^2 s}{q(1-hs)+1-q}\right]^n, \\
\text{i.e.} \quad & m \ln\left[\frac{q(1-s)+(1-q)(1-hs)}{q(1-hs)+1-q}\right] > n \ln\left[\frac{1-2q(1-q)hs-q^2 s}{q(1-hs)+1-q}\right], \\
\text{i.e.} \quad & m < n \frac{\ln\left[\frac{1-2q(1-q)hs-q^2 s}{q(1-hs)+1-q}\right]}{\ln\left[\frac{q(1-s)+(1-q)(1-hs)}{q(1-hs)+1-q}\right]} = n \times \beta_2(q,h,s).
\end{aligned}$$

Assuming $q \ll 1$ and $s \ll 1$, $\beta_1$ and $\beta_2$ can be approximated by $\beta_1 \approx q\frac{h}{1-h}$ and $\beta_2 \approx q$. This approximate value of $\beta_2$ is the same that the one defined by Nei et al. (1967) using similar assumptions, as expected. On autosomes, inversions should therefore increase in frequency when $m < nq$ and go to fixation when $m < \frac{nqh}{1-h}$. When permanently heterozygous (for instance if capturing the male-determining allele on the Y chromosome), inversions should go to fixation when $m < nq$, *i.e.* when they capture fewer mutations than the population average
